# A Single-Cell Atlas Reveals Cancer Cell Plasticity and Stromal-Immune Remodeling in Uterine Carcinosarcoma Across Diverse Ancestries

**DOI:** 10.64898/2026.04.21.720013

**Authors:** Santhilal Subhash, Marie-Thérèse Bammert, Brian Yueh, Timothy Chu, Mali Barbi, Aybuke Alici, Onur Eskiocak, Kadir A. Ozler, Aaron Nizam, Arielle Katcher, Charlie Chung, Vyom Shah, Elif Ozcelik, Nicolas Robine, Marina Frimer, Gary L. Goldberg, Semir Beyaz

## Abstract

Uterine carcinosarcoma (UCS) is an aggressive endometrial cancer defined by coexisting malignant epithelial and mesenchymal components, rapid metastatic dissemination, and poor therapeutic response. However, its cellular ecosystem remains poorly resolved, particularly in patients of African ancestry who are underrepresented in genomic datasets despite a disproportionate disease burden. Here, we generated a single-cell atlas of 15 primary and metastatic UCS specimens from a diverse cohort of 13 patients enriched for African ancestry, integrated with whole-genome sequencing. Malignant cells exhibited epithelial-like, mesenchymal-like, transitional, and stem/progenitor-like states within individual tumors that mapped to patient-specific copy number-defined subclones and RNA-velocity trajectories, supporting metaplastic state transitions. Compared to normal endometrium, primary tumors were enriched for epithelial-mesenchymal-transition (EMT), mTORC1, and glycolytic programs, whereas matched metastases show enhanced TNFα-NFκB-associated invasive programs. The tumor microenvironment contained immunosuppressive myeloid states and diverse cancer-associated fibroblast (CAF) subsets, including pericyte-like and matrix-remodeling subsets that act as predicted communication hubs through chemokine and immune-checkpoint circuits. We found a CAF-centered CCL2-CXCL1/2-IL10 module linked to CD8 T-cell dysfunction and a TIGIT-CD96-PVR checkpoint module. These data define the UCS cellular ecosystem in which malignant plasticity is coupled to stromal-immune cell remodeling in a patient cohort of enriched ancestries and nominate stromal-immune axes for further therapeutic investigation.

## Introduction

Uterine carcinosarcoma (UCS) is a rare but highly aggressive subtype of endometrial cancer that accounts for a disproportionate fraction of endometrial cancer mortality^1^. It is clinically characterized with high rates of metastasis, frequent recurrence and limited durable response to available therapies^1–6^. Women of African ancestry face a disproportionate burden, with higher incidence, more aggressive presentation, and lower five-year survival compared with women of European ancestry^7–9^. Despite this disparity, women of African ancestry remain markedly underrepresented in genomic and translational studies^10–15^. This gap has limited both biological insight and the development of therapeutic strategies that reflect the diversity of patients affected by this disease.

Histologically, UCS is composed of biphasic malignant epithelial and mesenchymal elements^16^. Genomic studies have shown that these components commonly share driver alterations, supporting a metaplastic carcinoma model in which phenotypic divergence emerges from a related malignant clone rather than from independent collision tumors^12,17,18^. Bulk genomic analyses, including TCGA-based studies and subsequent subtype classifications, have identified recurrent alterations in TP53, PI3K-pathway genes, and DNA-repair programs, and have implicated EMT-associated transcriptional programs and epigenetic remodeling in sarcomatous differentiation together with immunosuppressive myeloid programs^12,17–21^. However, bulk approaches cannot resolve how epithelial-like, mesenchymal-like, stem-like and transitional malignant states are organized within individual tumors, how these states relate to clonal architecture, or how stromal and immune compartments shape the UCS ecosystem.

Single-cell profiling provides a way to address these questions in cancers defined by histologic admixture and cellular plasticity^12,17,21,22^. In other aggressive tumors, malignant progression is shaped not only by tumor-intrinsic state transitions but also by coordinated stromal, myeloid, and T-cell programs that create niches for invasion, immune evasion, and therapeutic resistance^23–27^. Single-cell profiling in endometrial cancers more broadly has begun to define cancer-associated fibroblast (CAF) and exhausted T-cell programs, but did not focus on UCS or on ancestral diversity^28,29^. Whether similar ecosystem-level programs are present in UCS remains poorly understood, particularly in patients of diverse ancestries.

To address these gaps, we generated a single-cell atlas of primary and metastatic UCS integrated with matched whole-genome sequencing (WGS) in an ancestrally diverse cohort enriched for patients of African ancestry. We resolved patient-specific malignant states, copy number-defined subclones, immune phenotypes, and CAFs programs. Our data support a model in which UCS ecosystem reflects co-evolution of malignant epithelial-mesenchymal (E/M) plasticity with CAF-centered immune suppression and myeloid recruitment, nominating stromal-immune signaling axes for future therapeutic investigation.

## Results

### An ancestrally-enriched single-cell and genomic atlas resolves patient-specific UCS ecosystems

We profiled 15 UCS specimens from 13 patients, comprising 13 primary tumors and 2 matched metastatic lesions from the ovary and omentum (**Figure 1A**). The cohort was enriched for patients of African ancestry, a population that has been substantially underrepresented in prior UCS studies^12^. Whereas only 16 % of UCS cases in TCGA are from African-ancestry women, 62 % (8/13) of our patients self-identified as African ancestry and comprised diverse demographics for better representation (**Figure 1B**). We performed single-cell RNA sequencing (scRNA-seq) on all 15 tumors and whole-genome sequencing (WGS) on 12 tumors from 10 patients (**Figure 1C-F**). ADMIXTURE^30^ analysis of WGS data confirmed that our cohort is enriched for patients of African-ancestry, consistent with self-reported ancestry (**Figure 1C**, **Figure S1A**). MANTIS^31^ analysis identified one tumor (CS-014) as microsatellite-instability high (MSI-H), indicative of defective DNA mismatch repair, and associated with a high tumor mutational burden (TMB). Most tumors harbored TP53 alterations, reinforcing its enrichment in aggressive high grade uterine tumors^32,33^, together with recurrent mutations in the PI3K signaling pathway (e.g., PIK3CA, PTEN)^12,34,35^. Metastatic samples show overlapping but non-identical mutation patterns compared with primary tumors, indicative of clonal evolution during disease progression.

**Figure 1:**
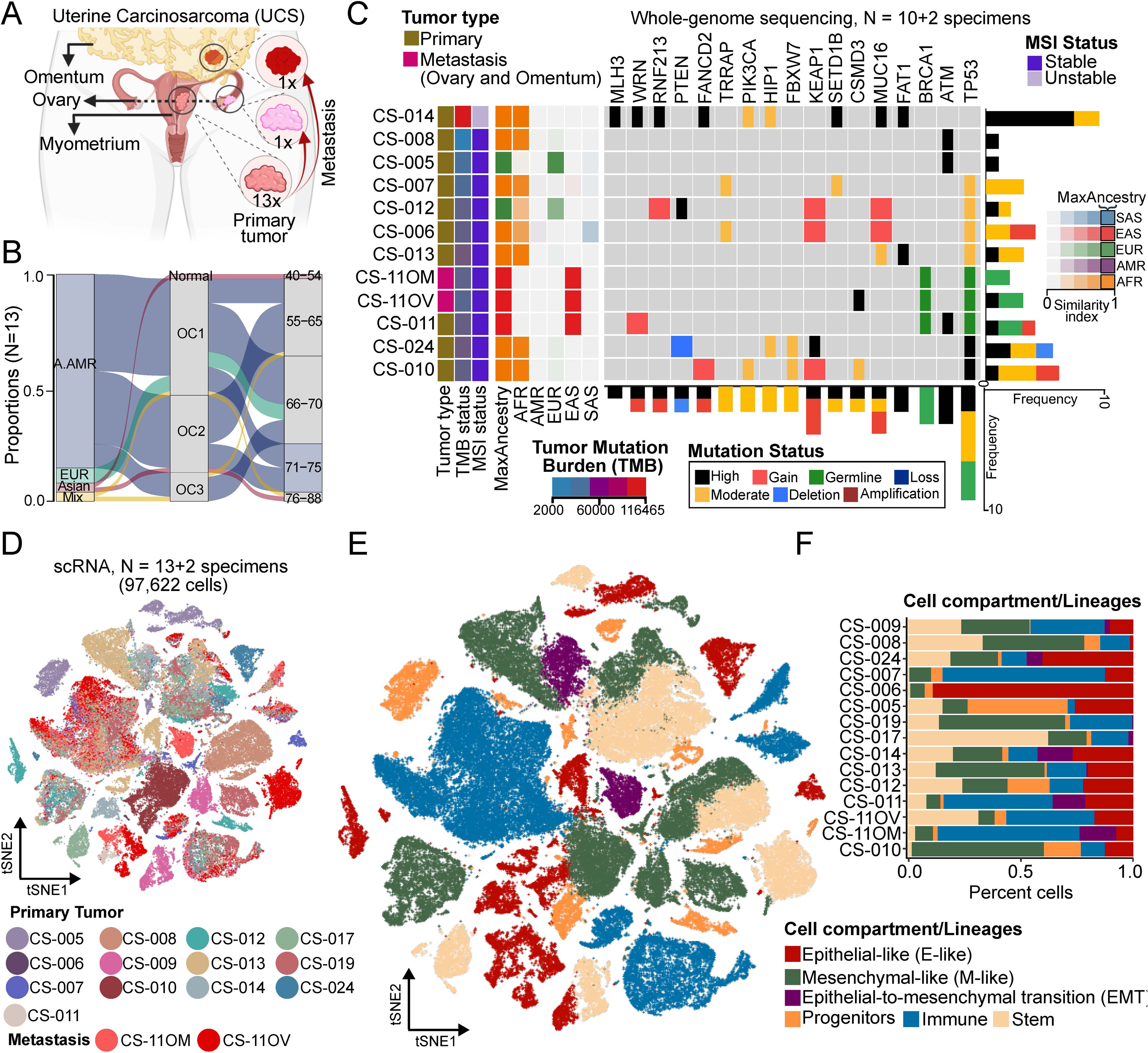
Ancestrally diverse single-cell transcriptome and genomic profiling of uterine carcinosarcoma reflects tumor heterogeneity. **(A)** Schematic overview of sample collection and tissue-of-origin. Tumor specimens were obtained from uterine carcinosarcoma (UCS) patients. Samples included primary uterine tumors (n = 13) and matched metastatic lesions from ovary and omentum (n = 2). Tissue was collected from anatomically distinct regions including endometrium, ovaries, and omentum. Downstream molecular profiling comprised single-cell RNA sequencing (scRNA-seq, n = 15 specimens) and bulk whole-genome sequencing (WGS, n = 12 specimens). **(B)** Alluvial plots shows the overall distribution of UCS patients (n = 13) included in this study according to self-reported ethnicity (m) n = 1, mixed ethnicity (MIX) n = 2), body mass index (BMI), and age groups. The BMI groups are classified based on their BMI indices such as Normal >= 18.5 & < 25; Obese class 1 (OC1) >= 25 & < 30; Obese class 2 (OC2) >= 30 & < 40; Obese class 3 (OC3) >= 40. **(C)** Oncoplot for WGS analysis of 10 primary UCS tumors and 2 matched metastases (Tumor type) showing tumor mutation burden (TMB status), microsatellite instability status (MSI status), and predicted ADMIXTURE-based ancestries. **(D)** Dimensionality reduction through t-SNE embedding of 97,622 cells derived from 13 primary tumors and 2 matched metastatic lesions (n = 15). Clusters in t-SNE are determined using 50 principal components with a resolution of 2.5. **(E)** t-SNE plot highlighting lineages of cells identified based on established gene expression markers. Annotated cell lineages include epithelial-like (E-like), mesenchymal-like (M-like), stem-like (Stem), epithelial-to-mesenchymal transition (EMT) states, progenitors, and immune populations. **(F)** Proportion bar plot shows the distribution of different cell lineages across individual UCS patients. X-axis denotes proportion or percentage of cells contributing to the cell lineages.

Across all tumors, we recovered 97,622 high-quality single cells and identified malignant and non-malignant compartments spanning epithelial-like (E-like), mesenchymal-like (M-like), EMT, stem-like/progenitor, stromal, and immune populations (**Figure 1D, E**). Cell type annotation leveraged canonical markers: E-like cells expressed *EPCAM*, *KRT19*, and *FOXJ1*; M-like cells expressed *ENO1*, *IGF2*, *PECAM1*, and *VWF*; stem/progenitor cells expressed *MKI67*, *PRC1*, *CD248*, and *MYC*; EMT cells expressed *CDH11*, *PRRX1*, *SNAI2*, and *ZEB1*; and immune cells expressed *PTPRC*/*CD45* (**Figure S1B**). The relative abundance of these lineages varied across tumors, highlighting inter-tumoral heterogeneity. Together, this ancestrally diverse single-cell atlas establishes a resource for UCS that captures the substantial patient-to-patient variability in malignant composition and tumor microenvironmental architecture.

### Copy number analysis reveals heterogeneous and plastic cancer cell states in UCS

To distinguish malignant from non-malignant cells while preserving ecosystem context, we inferred RNA-derived copy number alterations using CopyKAT (**Figure S2A**). This resulted in compartmentalization of aneuploid malignant cells from diploid stromal and immune populations, and a small fraction of ambiguous undefined cells, expressing stromal markers (*SPARC*, *MMP2*, *ACTA2*, *ENG*) and immune markers (*PTPRC*, *C1QA*, *IGKC*, *IL2RG*) (**Figure 2A**, **Figure S2A-B**). Transcriptional heterogeneity was greatest in the cancer compartment, intermediate in stromal cells, and lowest in immune cells (**Figure 2B**, **Figure S2C**). The malignant fraction consisted of E-like, M-like, EMT, and cancer stem cell-like (CSC-like) populations (**Figure 2C**), with their relative abundances varying substantially between tumors. Notably, CSC-like and E-like populations were consistently represented across UCS samples.

**Figure 2:**
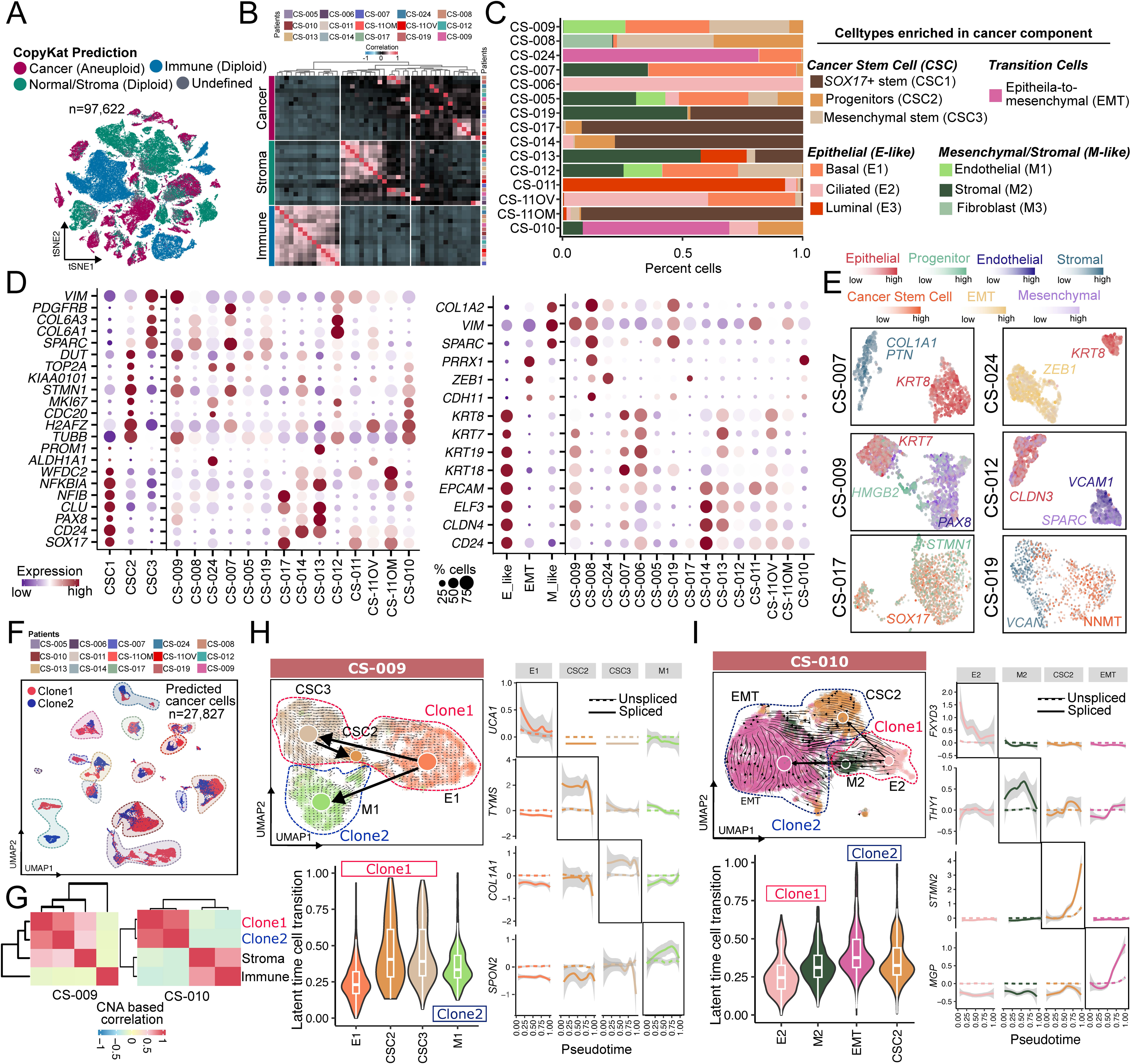
UCS cancer cells exhibit high level of heterogeneity, clonality, and cellular plasticity. **(A)** t-SNE embedding of single-cell transcriptomic data with cell populations classified using CopyKAT-based copy number inference, identifying aneuploid cancer cells, diploid stromal/normal cells, diploid immune cells, and unclassified cells. **(B)** Heatmap of pairwise Pearson correlations between UCS patients within cancer, stromal, and immune compartments, computed using the top 2,000 most variable genes per component identified by Seurat (VST). Color scale indicates correlation strength (red, positive; black, none; blue, negative). **(C)** Bar plot showing proportions of cancer cell states per patient, including stem-like (stem-progenitor, mesenchymal stem-like), epithelial (basal, ciliated, luminal), mesenchymal (endothelial, stromal, fibroblast-like), and EMT states. **(D)** Dot plots of marker gene expression defining stem-like, epithelial-like (E-like), mesenchymal-like (M-like), and EMT states across patients. Color intensity reflects expression level and dot size the fraction of expressing cells. Markers were identified using Seurat (log2FC ≥ ±1, adjusted p < 0.05, Wilcoxon test). **(E)** UMAP embedding of single-cell data per patient showing annotated cell populations and expression gradients of selected marker genes. **(F)** UMAP projection of cancer cells highlighting inferred clonal populations (n = 2 per patient) based on CopyKAT CNV profiles, with CNV segments clustered using Euclidean distance. **(G)** Heatmaps showing Pearson correlations between inferred cancer clones and other cellular compartments in representative patients (CS-009, CS-010), illustrating relationships between clonal identities and microenvironmental components. (**H**-**I**) UMAP projections of malignant cells from patients CS-009 (**H**) and CS-010 (**I**) with RNA velocity–derived trajectories overlaid on clonal identities. Violin plots show latent time distributions across cell states. Line plots depict gene-specific splicing dynamics along pseudotime (dotted, unspliced; solid, spliced; gray shading, standard deviation).

Within the malignant compartment, we identified multiple transcriptionally distinct subtypes. CSC-like cells segregated into three states: CSC1, CSC2, and CSC3 (**Figure 2C**). CSC1 expressed classical stem/progenitor-associated markers, including *SOX17*, *CLU*, *PAX8*, and *CD24* (**Figure 2D**). Notably, although *SOX17* acts as a tumor suppressor in non-gynecologic cancers, its elevated expression in gynecologic malignancies correlates with aggressiveness and metastatic potential^36^, and along with PAX8, reflects Müllerian lineage-specific uterine corpus endometrial carcinoma program^37^. CSC2 exhibited a proliferative, progenitor-like state marked by *CDC20*, *MKI67*, *STMN1*, and *KIAA0101*, a regulator of cancer progenitor growth and invasiveness in multiple tumor types^38,39^. CSC3 expressed mesenchymal stem-like genes, including *VIM* and *SPARC*. All CSC subtypes displayed highly heterogeneous expression patterns across UCS samples.

E-like cells resolved into basal (E1), ciliated (E2), and luminal-like (E3) states, each displaying substantial inter-tumoral variability (**Figure 2C**-**E**). EMT-like states were highly patient-specific and mostly present in CS-024 and CS-010 (**Figure 2C, E**). Together, these findings highlight that UCS cancer cells are profoundly heterogeneous, integrating diverse stem-like, E-like and M-like programs that are maintained within the tumor ecosystem.

### Copy number-defined subclones align with epithelial-mesenchymal state transitions

Ploidy-informed RNA-derived copy number profiles separated aneuploid malignant cells from diploid stromal and immune compartments and revealed multiple closely related malignant subclones within individual tumors (**Figure 2F**-**G**). Integrating these subclonal assignments with RNA velocity and latent-time analyses showed that genetically related malignant populations occupy distinct epithelial, stem/progenitor-like, EMT-like and mesenchymal transcriptional states. In a representative tumor, inferred trajectories progressed from epithelial-like or stem/progenitor-like states toward mesenchymal-like states, consistent with metaplastic state transitions (Figure 2H)^40^. Similar patterns were observed in additional UCS tumors (**Figure 2I**, **S2D**), where multiple subclones were anchored by distinct epithelial, progenitor, or mesenchymal programs. Because these analyses infer directionality from transcriptomic dynamics, we interpret them as supporting, rather than proving, a model in which UCS progression involves clonal diversification coupled to E/M plasticity.

### Distinct EMT and TNFα-NFkB programs are revealed in primary and metastatic UCS states

To delineate tumor-specific transcriptional programs, we compared UCS cancer cells with single-cell profiles from normal endometrium obtained from ten donors^41^. Donor cells, derived from women aged 18-34 years, were actively cycling, whereas UCS patients were primarily peri- or post-menopausal (40-88 years). To account for these cell cycle differences, we regressed normal endometrial cells for S- and G2M-phase signatures prior to comparison. This analysis identified 152 genes upregulated and 166 genes downregulated in UCS relative to normal tissue (**Figure 3A**). Primary UCS tumors showed broad upregulation of genes linked to malignant plasticity and stress adaptation, including *CLU*^42^, *LCN2*^43^, *S100A9*^44^, and *VIM*^45^. Pathway analysis identified enrichment of EMT, mTORC1 signaling, glycolysis, hypoxia, and unfolded protein response programs, consistent with a highly adaptive malignant state^46,47^. To investigate potential upstream regulators, we examined enriched transcription factor motifs among differentially expressed genes and identified cyclic-AMP-response element-binding protein (CREB)-associated motifs in upregulated genes (**Figure 3C**).

**Figure 3:**
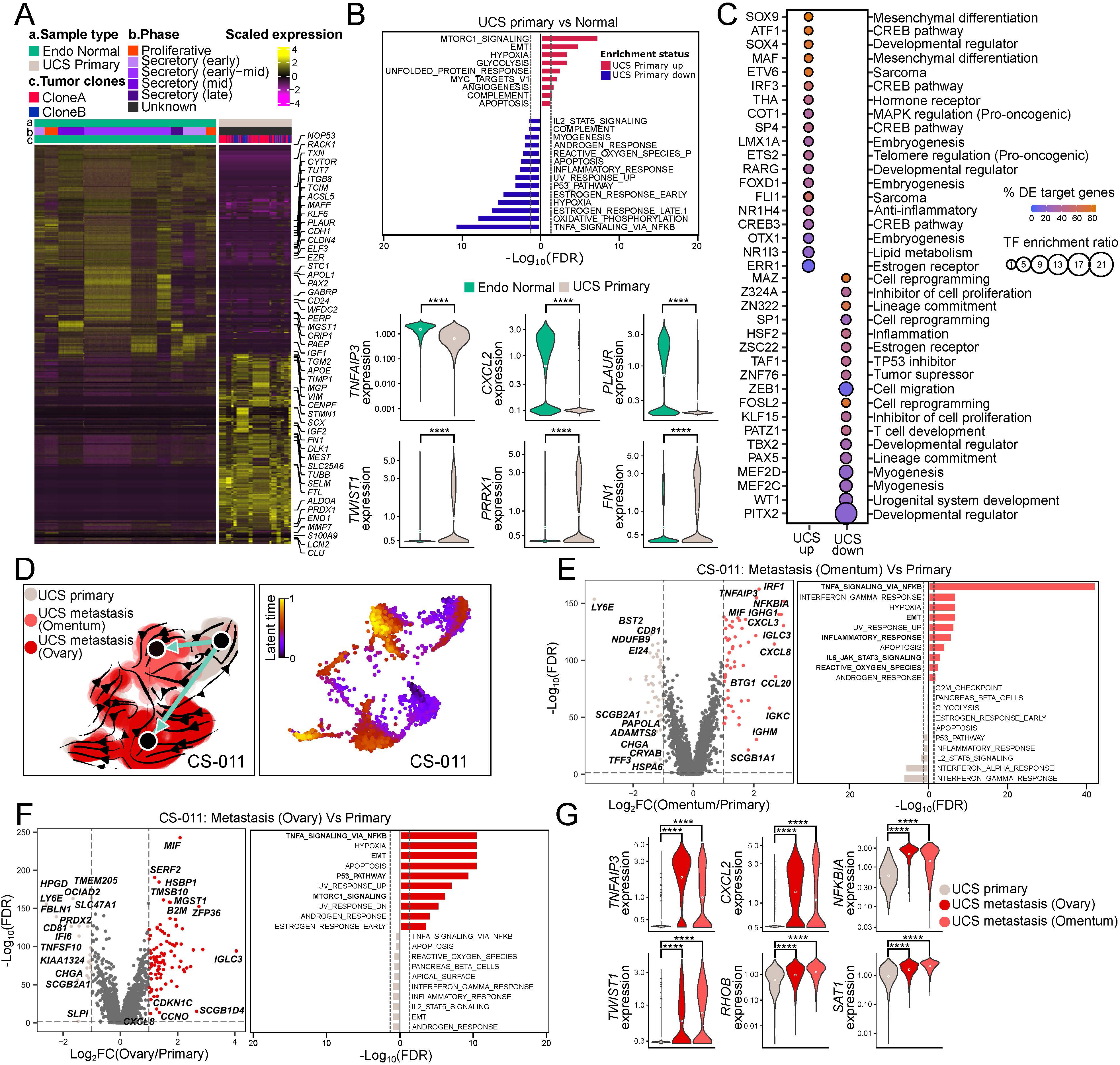
Trajectory-based analysis reveals molecular adaptations during metastatic progression. **(A)** Heatmap of top differentially expressed genes (p < 0.05, log2FC ±1) between normal endometrium and malignant clones from primary UCS tumors, identified using LIBRA and ranked by Wilcoxon rank-sum test. **(B)** Bar plots show enriched pathways among upregulated (red) and downregulated (blue) genes in UCS versus normal endometrium. Violin plots display expression of representative genes from TNF-α signaling (*TNFAIP3*, *CXCL2*, *PLAUR*) and EMT (*TWIST1*, *PRRX1*, *FN1*), with significance assessed by Wilcoxon test (ns ≥ 0.05; * < 0.05; ** < 0.01; *** < 0.001; **** < 0.0001). **(C)** Transcription factor motif enrichment in promoter regions (±250 bp from TSS) of differentially expressed genes (p < 0.05, log2FC ±1) in UCS versus normal endometrium. Color (violet–brown) indicates percentage of promoters with enriched motifs; dot size reflects enrichment ratio (MEME). **(D)** UMAP of single-cell profiles from primary tumor and matched ovarian and omental metastases (CS-011), showing inferred lineage trajectories (left) and latent time progression (right) computed using scVelo. **(E)** Volcano plot of differential expression between primary tumor and omental metastasis, with significant genes highlighted (log2FC ±1, adjusted p < 0.05). Bar plots show enriched pathways for upregulated (red) and downregulated (grey) genes. Analysis performed with LIBRA and GeneSCF. **(F)** Volcano plot of differential expression between primary tumor and ovarian metastasis with corresponding pathway enrichment (same thresholds and methods as in E). **(G)** Violin plots of TNF-α signaling (*TNFAIP3*, *CXCL2*, *NFKBIA*) and EMT-related genes (*TWIST1*, *RHOB*, *SAT1*) in UCS versus normal endometrium, highlighting inflammatory and mesenchymal reprogramming. Significance assessed using LIBRA (ns ≥ 0.05; * < 0.05; ** < 0.01; *** < 0.001; **** < 0.0001).

We next investigated metastatic differences by profiling two metastases (CS-011OM, omentum; CS-011OV, ovary) from the same patient. These metastases displayed distinct transcriptional landscapes and cell-cell communication networks compared with the matched primary tumor (CS-011) (**Figure 3D, Figure S3A**). RNA velocity and latent-time analyses revealed directional trajectories from primary to metastatic tumors, indicating sequential transcriptomic changes during tumor progression (**Figure 3D**). Compared with primary tumor, metastatic lesions exhibited pronounced TNF⍺-NFκB gene signatures coupled with EMT program engagement compared to the primary tumor (**Figure 3E-G, Figure S3B**), consistent with previous reports linking TNF⍺-driven EMT to metastatic competence^48–50^.

Together, these analyses indicate that UCS tumors exhibit cellular plasticity that associates with EMT, mTORC1 glycolytic and stress-adaptive programs, with TNF⍺-NFκB gene signatures coupled to invasive states in matched metastatic tumors.

### Tumor-infiltrating immune cells exhibit high inter-patient heterogeneity in UCS

Given the central role of the tumor microenvironment in shaping therapeutic response^51,52^, and the limited knowledge of immune landscapes in UCS, we profiled tumor-infiltrating immune cells (TIICs) across our cohort. TIICs comprised T-lymphocytes (CD4^+^, CD8^+^, and regulatory T-cells/Tregs), B-lymphocytes (B-cells, plasma B cells), myeloid populations (macrophages, monocytes, dendritic cells, mast cells), and innate lymphoid cells (ILCs) (**Figure 4A, S4A–B**) and their compositions varied across patients. For example, CS-005, CS-009, CS-012) were enriched for tumor-associated macrophages (TAMs), others (i.e., CS-007, CS-011) displayed high plasma B-cell infiltration (**Figure 4A**).

**Figure 4:**
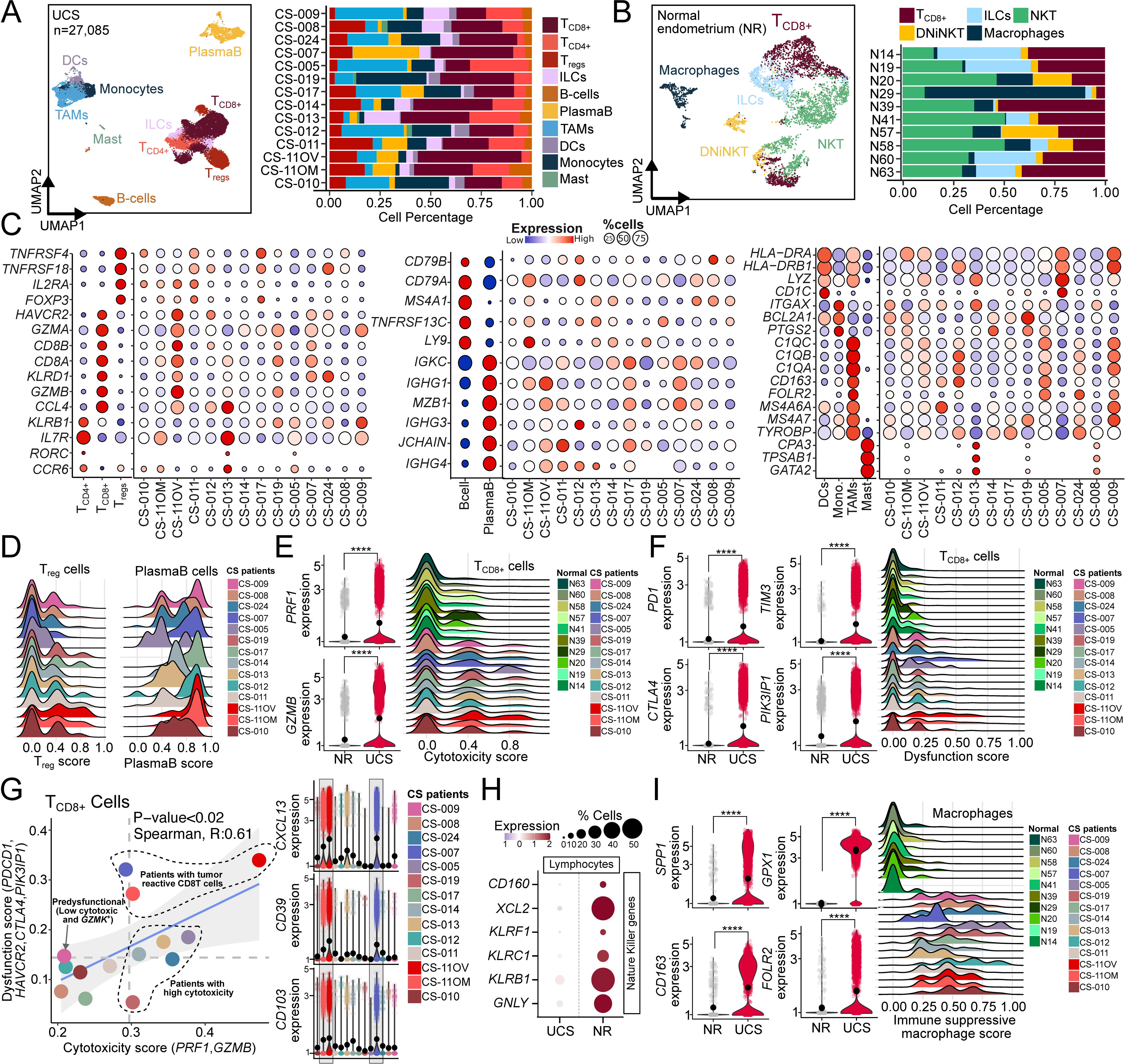
Integrated single-cell profiling reveals immune dysfunction and suppressive remodeling in UCS. **(A)** UMAP of single-cell transcriptomes highlighting annotated immune populations in UCS tumors, with bar plots showing relative proportions across patients. **(B)** UMAP of immune populations in normal endometrium (NR) with bar plots showing consistent cell-type proportions across donors, indicating a stable immune landscape compared to UCS. **(C)** Dot plots of lineage-specific marker expression across T cells (CD4^+^, CD8^+^, Tregs), B cells (B, plasma), and myeloid cells (DCs, monocytes, TAMs, mast cells) in UCS tumors. Dot size indicates fraction of expressing cells and color denotes expression level. Markers were identified by Seurat (log2FC ≥ ±1, adjusted p < 0.05, Wilcoxon test). **(D)** Ridge plots showing Treg (*FOXP3*, *IL2RA*) and plasma B cell (*JCHAIN*, *IGHG1*, *IGHG3*, *IGHG4*, *MZB1*) signature scores across UCS tumors, illustrating inter-patient variability. **(E)** Violin plots (left) show T-cell cytotoxic gene expression in NR and UCS. CD8^+^ cytotoxicity scores (*PRF1*, *GZMB*) are visualized by distribution and ridge plots (right). Significance assessed by LIBRA (ns ≥ 0.05; * < 0.05; ** < 0.01; *** < 0.001; **** < 0.0001). **(F)** Violin plots (left) show dysfunction/exhaustion gene expression in NR and UCS. CD8^+^ dysfunction scores (*PDCD1*, *HAVCR2*, *CTLA4*, *PIK3IP1*) are shown by distribution and ridge plots (right), with significance assessed by LIBRA. **(G)** Scatter plot shows Spearman correlation between cytotoxic and dysfunction signatures in CD8^+^ T cells from UCS. Violin plots show expression of tumor-reactive markers (*CXCL13*, *CD39*, *CD103*), highlighting functional heterogeneity. **(H)** Dot plot of classical NK cell marker expression across lymphocyte populations in UCS and normal tissues, with dot size indicating fraction of expressing cells and color denoting expression level. **(I)** Violin (left) and ridge plots (right) show expression and distribution of immunosuppressive macrophage signatures (*SPP1*, *FOLR2*, *CD163*, *GPX1*) in UCS versus normal endometrium, highlighting functional shifts. Significance assessed by LIBRA (ns ≥ 0.05; * < 0.05; ** < 0.01; *** < 0.001; **** < 0.0001).

By comparison, normal endometrium samples showed uniform immune profiles, with enrichment of T-lymphocytes, NK-cells, NKT cells and macrophages as the major myeloid population (**Figure 4B, Figure S4A-C**). The same immune cell clusters exhibited patient-specific transcriptional programs in UCS. For example, CD4^+^ T-cells expressing *IL7R*, *RORC*, *CCR6*, and *KLRB1* were restricted to certain UCS tumors (CS-005, CS-009, CS-010, CS-011, CS-024), whereas B-lymphocyte and myeloid subsets expressed distinct, tumor-specific markers (**Figure 4C**). Cell scoring revealed selective enrichment of T_regs_ (*FOXP3*, *IL2RA*) and plasma B cells (*JCHAIN*, *IGHG1-4*, *MZB1*) in individual UCS tumors (**Figure 4D**).

Cell-cell communication analysis predicted variable IL1 signaling during the progression from primary CS-011 to metastatic CS-011OM and CS-011OV (**Figure S4D**). IL1 signaling can promote cancer progression and metastasis through myeloid recruitment and *CXCR4* upregulation^53–55^, and we observed increased *CXCR4* expression in the metastatic lesions from this patient (**Figure S4D**). These results suggest that UCS tumors may leverage myeloid-mediated inflammation to sustain an aggressive microenvironment. Differential gene expression analysis between UCS and normal endometrium further showed transcriptional rewiring in both macrophage and T-cell compartments (**Figure S4E**).

### Distinct cytotoxic and immunosuppressive immune states are observed across UCS tumors

Building on the marked inter-patient heterogeneity in TIIC composition, we focused on CD8^+^ cytotoxic T-cells as central mediators of anti-tumor immunity and key predictors of immunotherapy response^56^. Several UCS tumors exhibited CD8^+^ T cells with expression of cytotoxicity genes (*PRF1*, *GZMB*)^57^ accompanied by dysfunction/exhaustion (*PDCD1*, *HAVCR2*, *CTLA4*, *PIK3IP1*)^58,59^, a phenotype absent in normal endometrium (**Figure 4E-G**). Notably, CD8⁺ T-cells from metastatic lesions (CS-011OM, CS-011OV) and one primary tumor (CS-007) contained particularly high cytotoxicity scores, suggesting patient-specific divergence of cytotoxic T cell responses. In parallel, UCS tumors showed a loss of NKT-cell signatures (C*XCL2*, *CD160*, *KLRF1*, *KLRC1*, *KLRB1*, *GNLY*)^60,61^ that are abundant in normal endometrium (**Figure 4H**), indicating a shift toward a CD8⁺-dominated cytotoxic compartment within the tumor microenvironment. Conversely, macrophages displayed an immunosuppression-associated gene expression profile^62^, marked by elevated *SPP1*, *CD163*, *FOLR2*, and *GPX1*^63–65^ (**Figure 4I**). Together, these results reveal the patient-specific cytotoxic and immunosuppressive immune states in UCS.

### UCS stroma is defined by abundant and diverse cancer-associated fibroblast subtypes

Cancer-associated fibroblasts (CAFs) are central stromal components in cancers, including UCS, where they orchestrate extracellular matrix (ECM) remodeling, regulate angiogenesis, and shape immune and metabolic tumor niches^66–68^. To explore the stromal cell landscape of UCS, we analyzed predicted stromal cell populations and found that ∼71 % of the UCS stromal cells were CAFs, ∼23 % were non-CAF populations, including stromal-like, epithelial-like, endothelial, and stem-like cells, and ∼6 % formed a mixed cluster expressing stromal, stem, and immune markers (**Figure S5A-B**). CAFs were stringently defined as CD90^+^/THY1^+^ fibroblasts lacking immune (PTPRC/CD45^-^) and endothelial (PECAM1^-^) signatures in our analysis (**Figure S5B-C**), while non-CAFs contained diverse subpopulations of endothelial, fibroblast, E-like, stem-like, and stromal signatures (**Figure S5D-E**).

CAFs can arise from diverse cellular origins, including endothelial cells, resident fibroblasts, epithelial cells, and bone-marrow-derived mesenchymal stem cells^69,70^. CD90^+^/THY1^+^ CAFs in our dataset segregated into five transcriptionally distinct subtypes (**Figure 5A-B**): pericyte-like CAFs (periCAFs), FAP^+^ CAFs, THBS4^+^ CAFs, antigen-presenting CAFs (apCAFs), and inflammatory CAFs (iCAFs)^71^. Quantification of CAF cell proportions revealed that most UCS tumors exhibited higher proportions of periCAFs and FAP^+^ CAFs, whereas THBS4^+^ CAFs, apCAFs, and iCAFs appeared in tumor-specific combinations (**Figure 5C**). In contrast, CAF subtype scoring, which reflects activation of subtype-specific gene expression programs rather than absolute cell abundance, demonstrated consistently high periCAF and apCAF signatures across UCS tumors (**Figure 5D**). Thus, although CAF populations vary in frequency, periCAF and apCAF transcription programs are broadly active across tumors, in line with their established roles in shaping immune-suppressive TME^72,73^. Multiple CAF subpopulations co-existed within individual tumors, and subtype-defining markers varied across samples, reflecting substantial molecular diversity (**Figure 5E**). For example, iCAFs in CS-019 uniquely expressed *CCL5*, whereas iCAFs in other tumors expressed *CXCR4* (**Figure 5E**).

**Figure 5:**
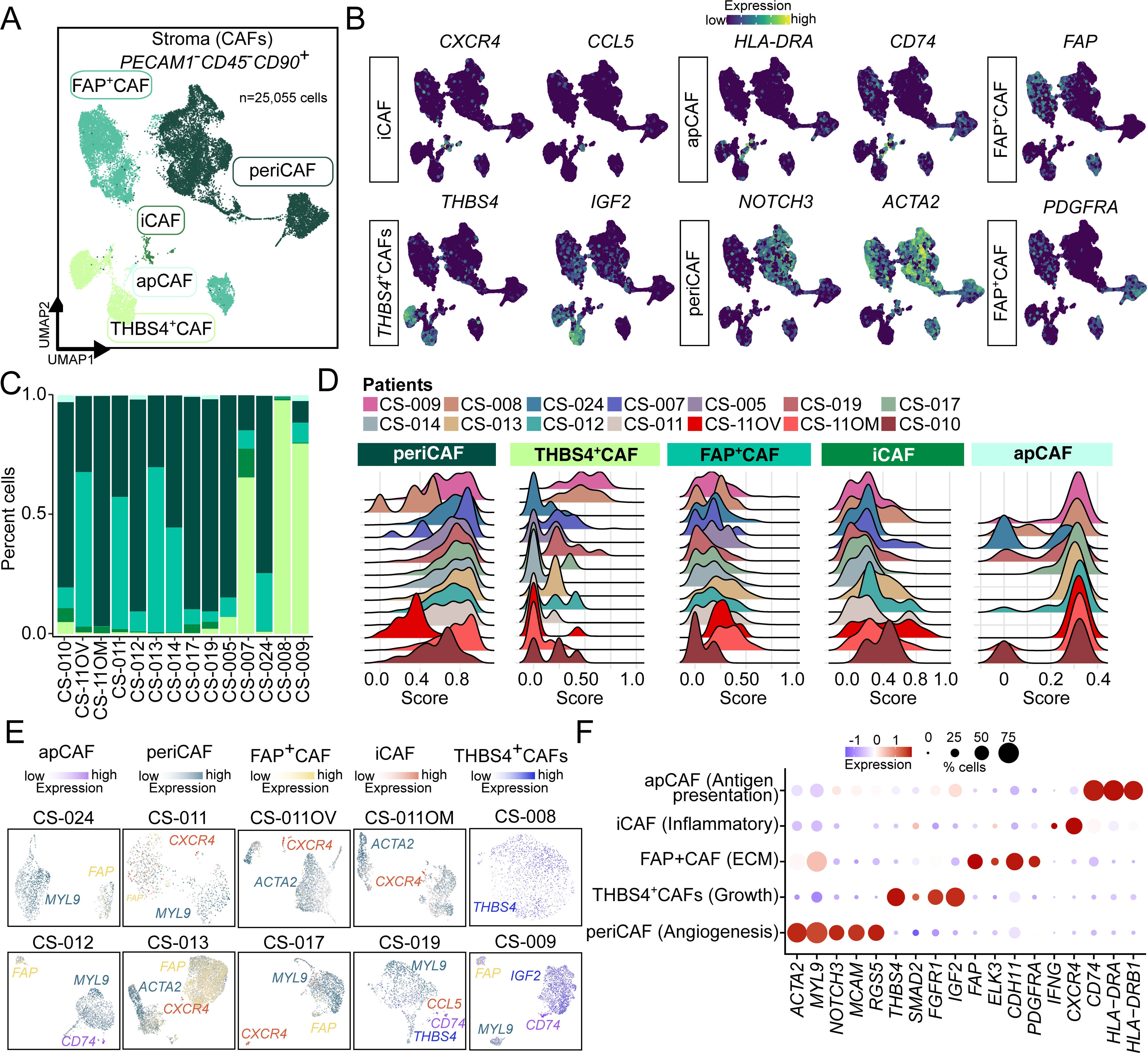
Distinct CAF subpopulations shape the stromal microenvironment in uterine carcinosarcoma. **(A)** UMAP of stromal fibroblasts from UCS tumors identifying distinct CAF subtypes, including inflammatory CAFs (iCAF), pericyte CAFs (periCAF), and antigen-presenting CAFs (apCAF), defined by marker expression and gating (*PECAM1*⁻, *CD45*⁻, *CD90*⁺). **(B)** UMAP feature plots showing expression of subtype-defining marker genes across CAF populations, supporting annotation of iCAF, periCAF, and apCAF states. **(C)** Bar plots of CAF subtype proportions across UCS patients, highlighting interpatient variability in stromal composition. **(D)** Ridge plots showing CAF signature scores across fibroblasts, including periCAF (*ACTA2*, *NOTCH3*), THBS4⁺ CAF (*THBS4*, *IGF2*), FAP⁺ CAF (*FAP*, *PDGFRA*), iCAF (*CXCR4*, *CCL5*), and apCAF (*CD74*, *HLA-DRA*). The x-axis denotes signature intensity and the y-axis the proportion of cells. **(E)** Patient-stratified UMAPs showing CAF subtype assignments (categorical colors) with overlaid marker gene expression (gradient), illustrating intra- and interpatient heterogeneity (n = 10). **(F)** Dot plot summarizing functional gene expression across CAF subtypes (e.g., ECM remodeling, inflammatory signaling, contractility/pericyte programs, antigen presentation). Dot size indicates fraction of expressing cells and color denotes expression level. Markers were identified by Seurat (log2FC ≥ ±1, adjusted p < 0.05, Wilcoxon test).

Each CAF subtype displayed specialized lineage-specific programs relevant to tumor progression (**Figure 5F**). PeriCAFs expressed pericyte and angiogenic regulators (*RGS5*, *MCAM*, *NOTCH3*), suggesting roles in vascular remodeling and increased vascular permeability that may facilitate leukocyte infiltration^74–76^. THBS4^+^ CAFs co-expressed growth factors (*FGFR1*, *SMAD2*, *IGF2*) implicated in tumor expansion across multiple cancers^77–79^ together with the ECM regulator *THBS4*, which has been linked to tissue remodeling and tumorigenesis^80,81^. FAP^+^ CAFs upregulated key ECM regulators (*ELK3*, *CDH11*, *PDGFRA*), consistent with a matrix-modifying role^82–84^. apCAFs exhibited robust MHC-II expression, supporting antigen-presenting potential^72,85^, whereas iCAFs expressed inflammatory mediators^86^ including *IFNG*, *CXCR4*, *CCL2*, *CXCL2*. These chemokines recruit leukocytes, including tumor-infiltrating myeloid cells (**Figure S5F**). Collectively, periCAFs and iCAFs expressed chemokine programs predicted to enhance myeloid recruitment, providing a potential link to the myeloid-rich, immune-suppressive tumor microenvironment^87^. These findings indicate that the UCS stromal cell compartment is not monolithic; instead, it contains distinct fibroblast states with likely non-redundant functions in vascular remodeling, matrix organization, immune regulation, and tumor support.

### CAF-directed chemokine and immune checkpoint circuits shape the UCS tumor immune microenvironment (TIME)

To map how stromal-immune interactions reinforce the tumor-supportive niche in UCS, we next mapped ligand-receptor interactions across stromal and immune compartments. This analysis revealed a dense network of CAF-centered signaling involving inflammatory (*IFN*, *IL*, *CCL*, *CXC*), immune checkpoint (*TIGIT*, *CD96*, *PVR*), suppressive (*BTLA*, *OX40*), and anti-tumor (*CD137*) pathways (**Figure 6A**), indicating that stromal-immune crosstalk is a major component of the UCS immune architecture. We selected significant cell communications based on patterns with high numbers of signaling genes having high contribution scores (>0.9). Among all cell types, periCAFs, FAP^+^CAFs, and CD8^+^ T cells exhibited the highest global communication activity (**Figure 6B**).

**Figure 6:**
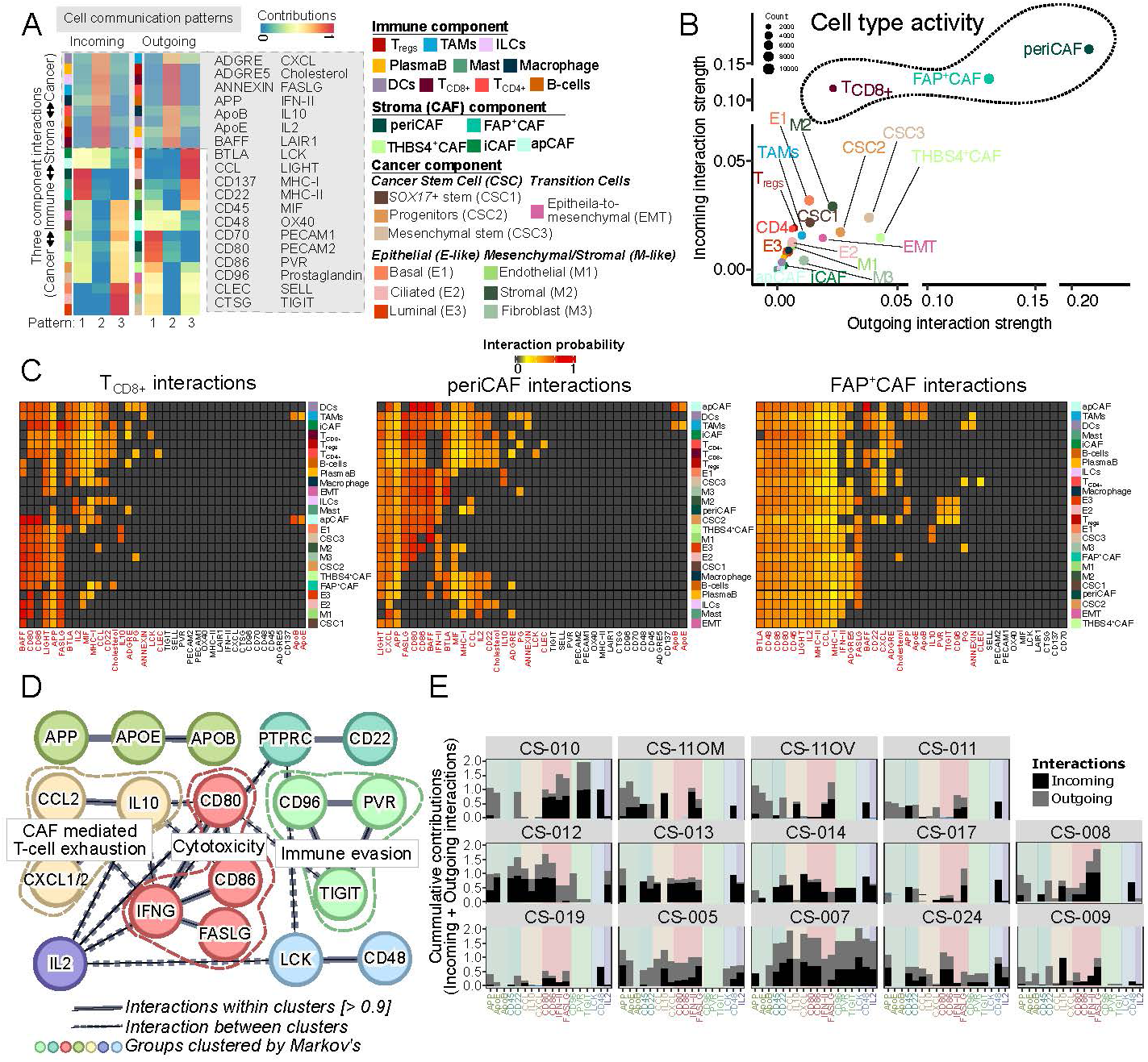
Compartment-specific cell-cell communication networks define stromal and immune signaling hubs in UCS. **(A)** Heatmap showing relative incoming and outgoing ligand–receptor signaling across Cancer, Stromal, and Immune compartments in UCS tumors. Pathways cluster into three compartment-dominant patterns, with top contributors highlighted. Color (blue–red) indicates low-to-high signaling. Significant interactions were identified by permutation testing against randomized cell group labels. **(B)** Scatter plot of outgoing versus incoming signaling strength across annotated cell types within each compartment. Highly active populations (e.g., CD8⁺ T cells, vCAFs, FAP⁺ CAFs) are highlighted. Signaling strength is defined by network centrality metrics (out-degree/in-degree) identifying dominant senders and receivers. **(C)** Heatmap of ligand-receptor interactions between high-activity populations (CD8⁺ T cells, vCAFs, FAP⁺ CAFs) and other tumor microenvironment cell types. Color intensity reflects interaction strength; red indicates statistically significant interactions. Interaction probabilities were estimated from ligand–receptor expression across cell types. **(D)** Protein-protein interaction network of signaling molecules derived from high-activity cell types, clustered using STRING (interaction confidence > 0.9) with the Markov Cluster Algorithm. Enriched modules are associated with immune regulation and exhaustion (PPI enrichment p < 1.0 x 10^-16^). Nodes represent proteins; edges denote functional/physical interactions, with thickness indicating confidence. **(E)** Bar plots of cumulative incoming and outgoing signaling strength per pathway across UCS patients, highlighting interpatient variability in tumor microenvironment communication dynamics.

Chemokine-cytokine networks are key drivers of tumor progression, enabling immune evasion and the recruitment of immunosuppressive populations, including T_regs_, TAMs, tumor-associated neutrophils (TANs), and myeloid-derived suppressor cells (MDSCs)^88^. These recruited leukocytes, particularly macrophages, dampen anti-tumor immunity and reinforce a suppressive microenvironment^89^. In UCS, FAP^+^CAFs prominently express CCL2 (MCP-1) and CXCL2, two chemokines linked to monocyte/macrophage recruitment^90^ (**Figure S5F**). TIME-wide ligand-receptor mapping further showed that CCL-CXCL signaling was largely confined to periCAFs and FAP⁺CAFs (**Figure 6C**). Through this axis, FAP^+^CAFs engaged both tumor-infiltrating immune cells and cancer cells. The CCL2-CXCL1/2 axis converged with IL10-mediated interactions (**Figure 4I**, **Figure 6D**), forming a CAF-dominated CCL2-CXCL1/2-IL10 module linked to exhausted CD8^+^ T cells. These data suggest a model in which CAF-derived chemokines contribute to T-cell dysfunction in UCS.

A second dominant signaling module centered on the TIGIT-CD96-PVR immune checkpoint axis (**Figure 6D**), which was highly specific to FAP⁺CAFs engagement with Tregs and cancer/epithelial cells. Given TIGIT’s established role in driving non-cytotoxic and suppressive T-cell states^91^, and the emerging involvement of PVR and CD96 in immune evasion^92^, this CAF-Treg-tumor triad suggests a stromal route for enforcing immune suppression in UCS. Notably, these signaling circuits varied across tumors (**Figure 6E**). The CCL2-CXCL1/2-IL10 axis was active in CS-005, CS-007, and CS-012, whereas the TIGIT-CD96-PVR module predominated in CS-007, CS-014, and CS-024, highlighting patient-specific immune microenvironmental states.

Together, these findings identify CAFs particularly periCAFs and FAP^+^CAFs as central components of the UCS TIME that are associated with chemokine-driven myeloid recruitment and immune checkpoint-mediated T-cell suppression. The CCL2-CXCL1/2-IL10 axis and the TIGIT-CD96-PVR axes therefore represent candidate stromal-immune entry points for therapeutic investigation in UCS.

## Discussion

UCS exemplifies extreme malignant plasticity, immune evasion, and therapeutic resistance, yet its ecosystem-level organization has remained poorly defined^18,21^. This study defines UCS as a co-evolving malignant-stromal-immune ecosystem shaped by E/M plasticity. By integrating scRNA-seq with WGS in an ancestry-enriched cohort, we resolve malignant cell states, inferred clonal architecture, immune dysfunction and CAF heterogeneity at individual patient resolution. Rather than collapsing inter-patient variability into a single consensus map, our analysis preserves patient-specific programs and reveals how distinct tumors combine E-like, M-like, stem-like and transitional malignant states with stromal and immune niches. This framework extends prior bulk genomic models of UCS by linking metaplastic malignant states to tumor microenvironmental remodeling.

Consistent with the biphasic histopathology of UCS^16,40^, malignant cells span epithelial, mesenchymal, and EMT-like states that coexist within individual tumors and across genetically-related clones. Several tumors display dominance of either E-like or M-like programs, whereas others harbor a continuum of states, underscoring pronounced inter- and intra-tumoral heterogeneity. EMT programs were prominent in matched metastatic lesions and in subsets of primary tumors, supporting a role for lineage plasticity in dissemination. Matched primary-to-metastatic analyses of the same patient suggests that metastatic competence may be encoded early and subsequently refined by microenvironmental selection, extending classical models of UCS histogenesis further^12,18,93^. The shared clonal architecture across epithelial, mesenchymal, and EMT states supports a conversion model, while the early diversification of malignant programs is also compatible with aspects of the combination model^12^. In contrast, we find little support for a pure collision model, as biphasic components are not molecularly independent. Together, these findings support a view of UCS as a dynamic continuum of malignant states shaped by ecosystem-level pressures.

At a single-cell level, UCS tumors exhibit extensive clonal heterogeneity, with distinct cancer cell subpopulations separated by inferred copy number alterations and transcriptional states. The variable expression of stem/progenitor-like developmental programs, such as *SOX17*^36,37,94^, further highlight molecular diversification within and between tumors. These findings underscore the need for future spatial characterization, single-cell WGS and functional studies to resolve clonal and cellular architecture of UCS.

The UCS TIME is profoundly reprogrammed and highly heterogeneous across patients. Compared with normal endometrium, UCS tumors exhibit enrichment of immunosuppressive myeloid populations and regulatory T cells, accompanied by loss of NK/NKT cell signatures. CD8^+^ T cells display variable cytotoxicity and exhaustion states, suggesting a spectrum of immune phenotypes ranging from tumor-reactive but dysfunctional to less activated states in UCS tumors This coordinated immune suppression profile provides a potential context for the limited efficacy of PD-1/PD-L1 blockade in UCS^95–97^ and emphasizes the importance of identifying alternative immune regulatory axes.

CAFs emerge as dominant organizers of the UCS ecosystem^98^. Multiple CAF states coexist within tumors, with periCAFs and FAP^+^ CAFs frequently acting as signaling hubs. CAF-centered ligand-receptor networks, particularly CCL2-CXCL1/2-IL10 interactions, are consistent with myeloid recruitment and T-cell dysfunction. In parallel, enrichment of the TIGIT-CD96-PVR immune-checkpoint axis highlights patient-specific mechanisms of immune evasion beyond canonical PD-1/PD-L1 signaling.

Several limitations should guide interpretation. First, although the cohort is substantial for a rare malignancy, the number of metastatic lesions is limited and derived from a single patient. Therefore, metastasis-associated programs should be verified in larger cohorts. Second, comparisons with normal endometrium may be affected by differences in donor age, hormonal state and tissue context despite cell-cycle regression. Third, RNA-derived copy-number inference, RNA velocity and ligand-receptor analyses infer clonal relationships, state trajectories and intercellular communication from transcriptomic data and should be validated with spatial and functional approaches. Future studies integrating spatial transcriptomics, multiplex imaging and perturbation models will be essential to test the CAF-immune and plasticity mechanisms nominated here.

Collectively, our data support a model in which UCS ecosystem emerges from the coupling of malignant lineage plasticity with stromal and immune reprogramming. The identification of patient-specific CAF-centered chemokine and checkpoint circuits provides a rationale for therapeutic strategies that combine tumor-intrinsic targeting with microenvironmental reprogramming. By resolving UCS biology in an ancestry-enriched cohort, this atlas provides a resource and conceptual framework for studying plastic, immune-suppressive and clinically aggressive carcinosarcomas.

## Methods

### Description of the Carcinosarcoma cohorts and Ethics statement

This study was approved by the Institutional Review Board at Northwell Health (IRB #18-0897) and Cold Spring Harbor Laboratory (IRB #21-17). All participants provided written informed consent, and all procedures were conducted in accordance with recognized ethical guidelines. Normal endometrium data used was obtained from previously published single cell study by Wang W et al^36^. All UCS patient metadata are summarized in **Table S1** including age, BMI, stage and ethnicity. Sex as a biological variable was not considered because this study is focused on endometrial cancer in female patients. Because sample sizes per ancestry are limited, we do not perform ancestry-stratified statistical comparisons.

### P1000 Carcinosarcoma cohort whole genome analysis

#### DNA extraction and library preparation

DNA was extracted from PBMCs isolated from patient blood or snap frozen tissue using the Zymo Quick-DNA Miniprep kit (Zymo, #D3024) following the manufacturer’s instructions. DNA quality and concentration were measured using a Nanodrop ND-1000 Spectrophotometer. Whole genome sequencing (WGS) libraries were prepared using the Truseq DNA PCR-free Library Preparation Kit (Illumina) in accordance with the manufacturer’s instructions. Briefly, 1 ug of DNA was sheared using a Covaris LE220 sonicator (adaptive focused acoustics). DNA fragments underwent bead-based size selection and were subsequently end-repaired, adenylated, and ligated to Illumina sequencing adapters. Final libraries were quantified using the Qubit Fluorometer (Life Technologies) or Spectromax M2 (Molecular Devices) and Fragment Analyzer (Advanced Analytical) or Agilent 2100 BioAnalyzer. Libraries were sequenced on an Illumina Novaseq6000 sequencer using 2×150bp cycles.

### Processing and analysis (Alignment, variant calling, Filtering, annotation)

#### Pre-processing

The New York Genome Center somatic pipeline (v6) was used to process and align the WGS data and call variants. Sequencing reads for the tumor and normal samples are aligned to the reference genome GRCh38 using BWA-MEM (v0.7.15) (arXiv:1303.3997v2 [q-bio.GN]). NYGC’s ShortAlignmentMarking (v2.1) is used to mark short reads as unaligned.

GATK (v4.1.0)^99^ FixMateInformation is run to verify and fix mate-pair information, followed by Novosort (v1.03.01) markDuplicates to merge individual lane BAM files into a single BAM file per sample. Duplicates are then sorted and marked, and GATK’s base quality score recalibration (BQSR) was performed. The result of the pre-processing pipeline is a coordinate sorted BAM file for each sample.

#### Quality control

Once preprocessing is complete, we compute several alignment quality metrics such as average coverage, %mapped reads and %duplicate reads using GATK (v4.1.0) and an autocorrelation metric (adapted for WGS from Zang et al.^100^) to check for unevenness of coverage. We also run Conpair^101^, a tool developed at NYGC to check the genetic concordance between the normal and the tumor sample and to estimate any inter-individual contamination in the samples.

#### Variant detection

The tumor and normal bam files are processed through NYGC’s variant calling pipeline which consists of MuTect2 (GATK v4.0.5.1)^102^, Strelka2 (v2.9.3)^103^ and Lancet (v1.0.7)^104^ for calling Single Nucleotide Variants (SNVs) and short Insertion-or-Deletion (Indels), SvABA (v0.2.1)^105^ for calling Indels and Structural variants (SVs), Manta (v1.4.0)^106^ and Lumpy (v0.2.13)^107^ for calling SVs and BIC-Seq2 (v0.2.6)^108^ for calling Copy-number variants (CNVs). Manta also outputs a candidate set of Indels which is provided as input to Strelka2 (following the developers recommendation, as it improves Strelka2’s sensitivity for calling indels >20nt).

#### Variant merging

Next, the calls are merged by variant type (SNVs, Multi Nucleotide Variants (MNVs), Indels and SVs). MuTect2 and Lancet call MNVs, however Strelka2 does not, and it also does not provide any phasing information. So to merge such variants across callers, we first split the MNVs called by MuTect2 and Lancet to SNVs, and then merge the SNV callsets across the different callers. If the caller support for each SNV in a MNV is the same, we merge them back to MNVs. Otherwise those are represented as individual SNVs in the final callset. Lancet and MantaSV are the only tools that can call deletion-insertion (delins or COMPLEX) events. Other tools may represent the same event as separate yet adjacent indel and/or SNV variants. Such events are relatively less frequent, and difficult to merge. We therefore do not merge COMPLEX calls with SNVs and Indels calls from other callers. The SVs are converted to bedpe format, all SVs below 500bp are excluded and the rest are merged across callers using bedtools^109^ pairtopair (slop of 300bp, same strand orientation, and 50% reciprocal overlap).

#### Somatic variant annotation (SNVs, Indels, CNVs, and SVs)

SNVs and Indels are annotated with Ensembl as well as databases such as COSMIC (v86)^110^, 1000Genomes (Phase3)^111^, ClinVar (201706)^112^, PolyPhen (v2.2.2)^113^, SIFT (v5.2.2)^114^, FATHMM (v2.1)^115^, gnomAD (r2.0.1)^116^ and dbSNP (v150)^117^ using Variant Effect Predictor (v93.2)^118^. For CNVs, segments with log2 > 0.2 are categorized as amplifications, and segments with log2 < -0.235 are categorized as deletions (corresponding to a single copy change at 30% purity in a diploid genome, or a 15% Variant Allele Fraction). CNVs of size less than 20Mb are denoted as focal and the rest are considered large-scale.

We use bedtools^109^ for annotating SVs and CNVs. All predicted CNVs are annotated with germline variants by overlapping with known variants in 1000 Genomes and Database of Genomic Variants (DGV) (22). Cancer-specific annotation includes overlap with genes from Ensembl^119^ and Cancer Gene Census in COSMIC, and potential effect on gene structure (e.g. disruptive, intronic, intergenic). If a predicted SV disrupts two genes and strand orientations are compatible, the SV is annotated as a putative gene fusion candidate. Note that we do not check reading frame at this point. Further annotations include sequence features within breakpoint flanking regions, e.g. mappability, simple repeat content and segmental duplications.

### Somatic variant filtering

#### Panel Of Normals

The Panel Of Normals (PON) filtering removes recurrent technical artifacts from the somatic variant callset^102^.

#### PON generation

The Panel of Normals for SNVs, indels and SVs was created with whole-genome sequencing data from normal samples from 242 unrelated individuals. Of these, sequencing data for 148 individuals was obtained from the Illumina Polaris project which was sequenced on the HiSeqX2 platform with PCR-free sample preparation. The remaining samples were sequenced by the NYGC. Of these, 73 individuals were sequenced on HiSeqX, 11 on NovaSeq, and 10 were sequenced on both. We ran MuTect2 in artifact detection mode and Lumpy in single sample mode on these samples. For SNVs and indels, we created a PON list file with sites that were seen in two or more individuals.

For SVs, we used SURVIVOR (v1.0.3)^120^ to merge Lumpy calls. Variants were merged if they were of the same type, had the same strand orientation, and were within 300bp of each other (maximum distance). We did not specify a minimum size. After merging SVs, we used these calls as a PON list.

#### PON filtering

For SNVs and Indels, we use the PON list to filter the somatic variants in the merged SNV and indel files. To filter our somatic SV callset, we merge our PON list with our callset using bedtools pairtopair (slop of 300bp, same strand orientation, and 50% reciprocal overlap), and filtered those SVs found in two or more individuals in our PON.

#### Common germline variants

In addition to the PON filtering, we remove SNVs and Indels that have minor allele frequency (MAF) of 1% or higher in either 1000Genomes (phase 3) or gnomAD (r2.0.1)^116^, and SVs that overlap DGV and 1000Genomes (phase3). CNVs are annotated with DGV and 1000 Genomes but not filtered.

#### All Somatic and High-confidence variants

Variants that pass all of the above-mentioned filters are included in our final somatic callset (hereby referred to as AllSomatic). For SNVs, indels and SVs, we also annotate a subset of the somatic callset as high confidence. For SNVs and indels, high confidence calls are defined as those that are either called by two or more variant callers, or called by one caller and also seen in the Lancet validation calls or in the Manta SV calls.

For structural variants, high confidence calls are taken from the somatic callset if they meet the following criteria: called by 2 or more variant callers, or called by Manta or Lumpy with either additional support from nearby CNV changepoint or split-read support from SplazerS^116^, an independent tool used to calculate the number of split-reads supporting SV breakpoints. An SV is considered supported by SplazerS if it found at least 3 split-reads in the tumor only. Nearby CNV changepoints are determined by overlapping BIC-Seq2 calls with the SV callset using bedtools closest. An SV is considered to be supported by a CNV changepoint if the breakpoint of the CNV is within 1000bp of an SV breakpoint.

### MSI detection

We run MANTIS (v1.0.4)^31^ for Microsatellite Instability (MSI) detection in microsatellite loci (found using RepeatFinder, a tool included with MANTIS). A sample is considered to be microsatellite unstable if it’s Step-Wise Difference score reported by MANTIS is greater than 0.4 (or 0.62 in absence of a matched-normal). Otherwise, it is considered to be microsatellite stable (MSS).

### Genetic ancestry estimation

Ancestry proportion is determined by the software ADMIXTURE^30^ v1.3.0, which uses a maximum likelihood-based method to estimate the proportion of reference population ancestries in a sample. We genotyped the reference markers generated from 1,964 unrelated 1000 Genomes project samples directly on the samples using GATK pileup. Individuals from populations MXL (Mexican Ancestry from Los Angeles USA), ACB (African Caribbean in Barbados), and ASW (African Ancestry in Southwest US) were excluded from the reference due to being putatively admixed. The reference was further filtered by using only SNP markers with a minimum minor allele frequency (MAF) of 0.01 overall and 0.05 in at least one 1000 genomes superpopulation. Variants are additionally linkage disequilibrium (LD) pruned using PLINK v1.9 with a window size of 500kb, a step size of 250kb and r2 threshold of 0.2. The analysis results in a proportional breakdown of each sample into 5 continental populations (AFR, AMR, EAS, EUR, SAS) and 23 sub-populations.

### Single Cell Profiling by Chromium 10X sequencing and Analysis

#### Tissue processing, cell preparation, and library construction

Surgical resections of 15 specimens (13 primary and 2 metastasis) from 13 UCS patients were taken and the tissues were dissociated according to Katcher et al.^121,122^. In brief, tissue was washed with cold 1X PBS and minced into small (∼0.25 cm) fragments, before being transferred to 5 ml of RPMI media (Sigma Aldrich, #R8758) containing 1 mg/mL Collagenase (Sigma Aldrich, #C9407) and 10 µM Y-27632 dihydrochloride (Tocris, #1254). Tissue suspension was transferred to a rotating incubator (speed: 30 rpm, 37 °C) for 60-120 min depending on successful tissue dissociation. Supernatant was transferred and centrifuged at 300 g for 5 min at room temperature (RT). Supernatant was discarded and the pellet was resuspended in 3 mL of TrpLE Express Enzyme (1X, no phenol red, Thermo Fisher Scientific, #12604103) with 10 µM Y-27632 dihydrochloride followed by an incubation of 10-20 min at 37 °C with regular mixing. To stop the reacting 3 mL of ADMEM/F12 (Life Technologies, #312634028) was added and the cell suspension centrifuged at 300 g for 5 min at RT. The sample was then filtered through a 40um cell strainer (Corning, #431750) and dead cell removal was conducted using the Dead Cell Removal Kit (Miltenyi Biotec 130-090-101), following the manufacturer’s instructions. Samples averaged 80-90% viability with at least 1.5 million cells. Samples are processed for sequencing with Chromium Next GEM Chip G (10x Genomics). GEM generation and barcoding, reverse transcription, cDNA generation and library construction using 3ʹ Gene Expression Library Construction using the Chromium Single Cell 30 Library (v3 chemistry) following the manufacturer’s protocol. Dual-indexed, single-cell libraries were pooled and sequenced in paired-end reads on Novaseq (Illumina).

#### Single-cell analysis

Raw reads from FASTQ files are aligned to reference genome GRCh38 and quantified for GRCh38.84 gene annotation using CellRanger (v.6.0.0) (https://10xgenomics.com). First, the CellRanger *mkfastq* command with the CellRanger sample sheet was used to demultiplex the base call files for each flow cell into FASTQ files. Second, the CellRanger *count* command was called to generate single cell feature counts for each library by specifying the library name in the argument. The filtered feature barcode matrix was used for further data analysis.

#### Quality check and clustering

The downstream analysis using feature barcode matrix was performed using Seurat (v.3.0.0) package. Individual patients with their feature barcode matrix libraries were converted into Seurat object using *Read10X* and *CreateSeuratObject*. For each individual patients we removed cells with high mitochondrial content (>20%) and cells with less than 200 genes. These cut-offs are based on QC inspection and previous single cell studies on endometrium/ovarian^41,123^. Subsequently, the gene in remaining 97,622 cells were log normalized, variable genes detected, scaled and the principal components computed. The top principal components were identified and used for the UMAP and t-SNE dimensionality reduction. Normal endometrium singe cell data from *Wang W et al*.^41^, was filtered using the cutoffs mentioned in the publication and the clustering is performed like UCS patients.

#### Heterogeneity and Assignment of cell types

Initial clustering was performed, and UCS patients showed highly heterogenous profiles. Individual clusters are assigned to a patient based on proportion of cells from the patient contributing to the cluster. Hence the gene markers are predicted per cluster in individual UCS patient by *FindAllMarkers* function from Seurat. This method of assigning gene markers and cell types preserved both molecular and cellular-level heterogeneity in UCS patients. Similar approach was used in prediction of gene markers and cell type assignment for normal endometrium singe cell data from *Wang W et al*.^41^ Scoring of cells for different functional modules are done using *AddModuleScore_UCell* function from UCell R package.

### RNA abundance-based copy number alteration (rCNA) profiling

Later the cells from UCS patients were subjected to CopyKat analysis with default parameters to determine cancerous and non-cancerous cells in patients’ single-cell data by extracting ploidy information by measuring RNA copy number abundance (rCNA) in the genome (https://github.com/navinlabcode/copykat)^124^. The Aneuploid cells were assigned cancerous and remaining diploid cells as non-cancerous. We used hierarchical clustering with Euclidean distance measure to segregate cancerous cells from non-cancerous cells and to determine subclonal profiles within predicted cancer cells. We tested the subclonal profiles using different cutoffs for number of trees (subclusters) and found 2 optimal tree (subcluster) per patient(https://github.com/decodebiology/Carcinosarcoma_singlecell_atlas). We were able to separate non-cancerous diploid cells as immune and non-immune (Stroma) profiles based on the gene markers (**Figure S2B**). The undefined cells by CopyKat express both stroma and immune gene markers and was excluded from further analysis. The individual components such as cancer, immune, and stroma was further analyzed separately. We have performed ligand-receptor interaction analysis to identify cell-cell communications and signaling in the individual components using CellChat^125^.

### Differential expression analysis

Differential expression (DE) analysis between normal endometrium and UCS predicted cancer cells were performed by implementing Libra using *run_de* function (https://github.com/neurorestore/Libra)^126^. We used single cell appropriate statistical method Wilcoxon Rank-Sum test to find differentially expressed genes. Similarly, DE analysis was performed between UCS primary and metastatic stages, between normal endometrium immune profile and UCS tumor infiltrating immune profile. Significant DE genes were selected based on adjusted p-value (<0.05) and log-fold-change (±1.0) cutoffs. Obtained DE genes were subjected to functional enrichment analysis with human MSigDB hallmark gene sets (https://gsea-msigdb.org) and gene ontology (https://geneontology.org) using GeneSCF (v1.1-p3) tool^127^. The enriched gene sets are selected by p-value<0.05 cutoff.

### Promoter Sequence Based Motif Enrichment Analysis

Using custom scripts we extracted nucleotide sequences (FASTA) from the promoters (± 250 bp from TSS) of list of significantly differentially expressed genes between UCS predicted cancer cells and normal endometrium (https://github.com/decodebiology/extract_promoter_sequence) with GRCh38.84 gene and genome annotation. Obtained FASTA sequences were used to find enriched transcription factor motifs using SEA (v5.5.5) from MEME tool^128^. Motif database HOCOMOCO (v11) full human was used, and the shuffled primary sequences preserving 3-mer frequencies are considered as control sequences. Also, 10% of the input sequences were randomly assigned to the hold-out set to improve p-value accuracy. The motifs are selected based on enrichment E-value of 10 or smaller. Motifs analysis was performed independently on both up and downregulated genes.

### Cell-Cell Communication Analysis by Ligand-receptor Interactions

We have performed ligand-receptor interaction analysis to identify cell-cell communications between three components of UCS tumors such as cancer-immune-stroma/CAF using CellChat^125^. From the analysis we obtained multiple patterns of interaction profiles between cell types and individual patterns are clustered based on the interaction strength between cell types. We selected significant cell communications based on patterns with a greater number of cell types interacting with high contribution score (>0.9). Signaling (ligand-receptor interactions) that are contributing to this cell-cell communications between these cell types are extracted for further analysis. Cell types having higher activity was assigned using incoming and outgoing signal strengths. We looked for other cell types that are interacting with highly active cell types with the corresponding signaling contributions resulting in interactions. These signaling genes were then subjected to StringDB to find coregulation patterns among them.

## Data and Code Availability

The genomic sequencing data will be deposited in dbGAP repository with accession and the single cell RNA-seq dataset will be accessed from GEO repository upon acceptance of the manuscript. All the pipelines and the code used for the analysis has been deposited in github, https://github.com/decodebiology/Carcinosarcoma_singlecell_atlas. The single-cell dataset generated and/or analyzed in this study can be explored through an interactive portal available at: https://decodebiology.shinyapps.io/UCS_Tumor_Tissue/.

## Acknowledgements

We thank the members of the Beyaz lab for critical discussions. We thank Northwell Health Biospecimen Repository and the Gynecologic Oncology staff for assistance in recruitment of patients from diverse demographics and acquisition of specimens for this study. We thank Cold Spring Harbor Laboratory Cancer Center Shared Resources (Flow Cytometry, Microscopy, Sequencing, Organoid and Histology Core Facilities) supported in part by the National Cancer Institute Cancer Center Support Grant (5P30CA045508). This work was financially supported by grants to S.B from the National Cancer Institute (R37CA292807), Oliver S. and Jennie R. Donaldson Charitable Trust, the Mark Foundation for Cancer Research (20-028-EDV), the Beverly Hazelkorn Uterine Cancer Research Foundation, the Cold Spring Harbor Laboratory and Northwell Health Affiliation, the Foundation for Women’s Health, and the New York Genome Center Polyethnic-1000 Initiative. The sample processing, computations, and depositions of the datasets from this study and TCGA was performed on High Performance Computing (HPC) resources Elzar and Agastya provided by Cold Spring Harbor Laboratory and Indian Institute of Technology Jammu respectively.

