## Supplemental for "A Single-Cell Atlas Reveals Cancer Cell Plasticity and Stromal-Immune Remodeling in Uterine Carcinosarcoma Across Diverse Ancestries": 20260702-Supplemental Figures + figure legends.pdf

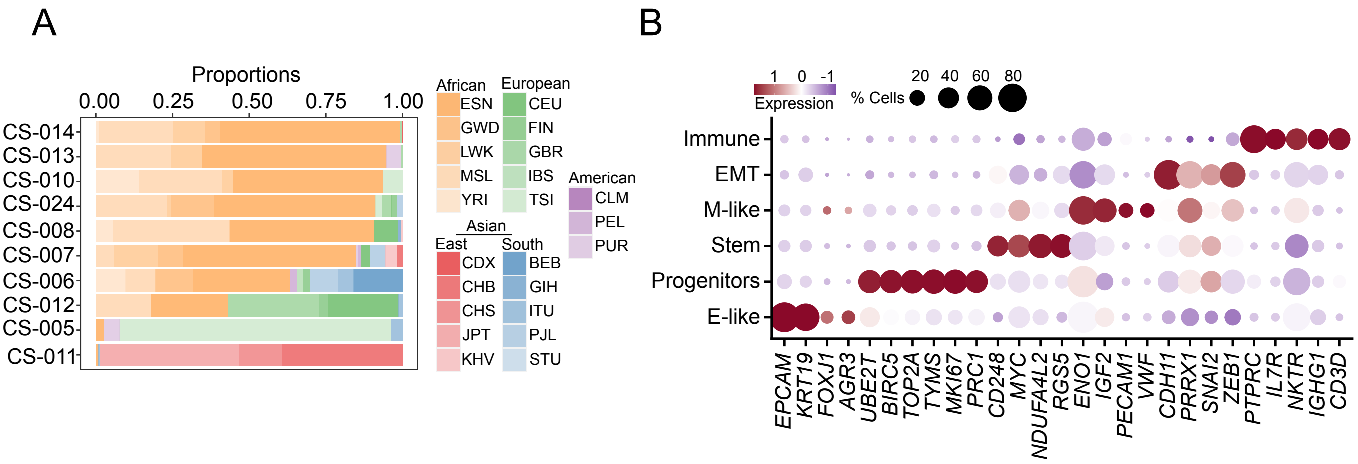
A
Proportions
0.00
0.25
0.50
0.75
1.00
CS-014
CS-013
CS-010
CS-024
CS-008
CS-007
CS-006
CS-012
CS-005
CS-011
African
ESN
GWD
LWK
MSL
YRI
European
CEU
FIN
GBR
IBS
TSI
American
CLM
PEL
PUR
Asian
East
South
CDX
CHB
CHS
JPT
KHV
BEB
GIH
ITU
PJL
STU
B
1 0 -1
Expression
% Cells
20 40 60 80
Immune
EMT
M-like
Stem
Progenitors
E-like
EPCAM
KRT19
FOXJ1
AGR3
UBE2T
BIRC5
TOP2A
TYMS
MKI67
PRC1
CD248
MYC
NDUFA4L2
RGS5
ENO1
IGF2
PECAM1
VWF
CDH11
PRRX1
SNAI2
ZEB1
PTPRC
IL7R
NKTR
IGHG1
CD3D

**Figure S1:**

(**A**) ADMIXTURE-based ancestry inference demonstrates diverse continental representation across the cohort, with clear assignment to major ancestral subgroups. ESN: Esan (Nigeria), GWD: Gambian in Western Division (Gambia), LWK: Luhya in Webuye (Kenya), MSL: Mende in Sierra Leone, YRI: Yoruba in Ibadan (Nigeria), CEU: Utah Residents, FIN: Finnish, GBR: British, IBS: Iberian (Spain), TSI: Toscani (Italy), CDX: Chinese Dai in Xishuangbanna, CHB: Han Chinese in Beijing, CHS: Southern Han Chinese, JPT: Japanese in Tokyo, KHV: Kinh in Ho Chi Minh City (Vietnam), BEB: Bengali from Bangladesh, GIH: Gujarati from Houston, ITU: Indian Telugu from the UK, PJL: Punjabi from Lahore (Pakistan), STU: Sri Lankan Tamil, CLM: Colombians from Medellin, PEL: Peruvians from Lima, PUR: Puerto Ricans.

(**B**) Dot plot of canonical marker genes used to define cellular compartments or lineages. Dot size indicates the proportion of expressing cells, and color intensity reflects average expression level (red, high; blue, low; white, undetected), supporting robust and distinct cell-type classification.

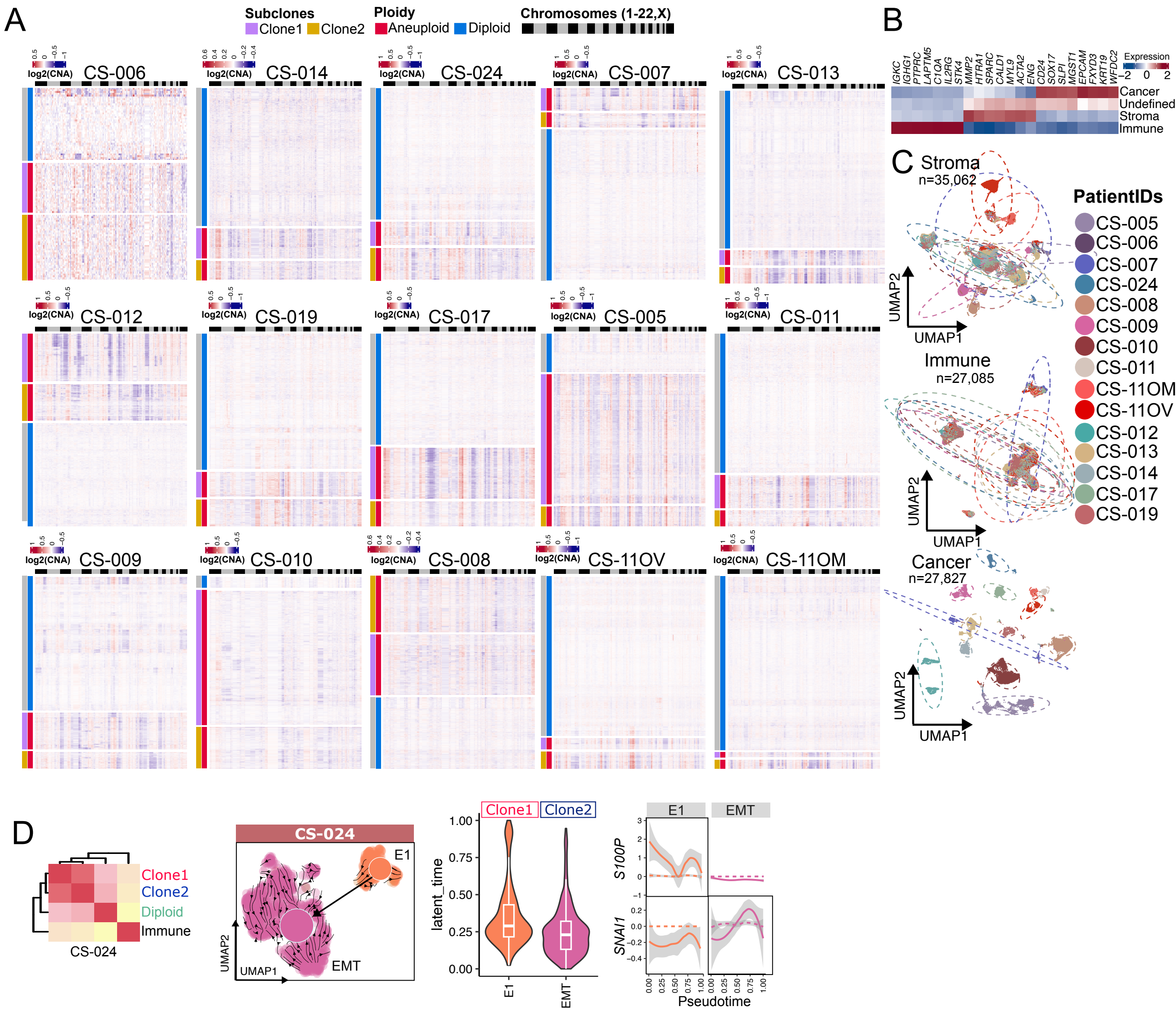

A
Subclones
Clone1 Clone2
Ploidy
Aneuploid Diploid
Chromosomes (1-22,X)
log2(CNA)
CS-006
CS-014
CS-024
CS-007
CS-013
CS-012
CS-019
CS-017
CS-005
CS-011
CS-009
CS-010
CS-008
CS-11OV
CS-11OM
B
IGKC IGHG1 PTPRC LAPTM5 C1QA IL2RG STK4 MMP2 HTRA1 SPARC CALD1 MYL9 ACTA2 ENG CD24 SOX17 SLPI MGST1 EPCAM FXYD3 KRT19 WFDC2
Expression
Cancer
Undefined
Stroma
Immune
C
Stroma
n=35,062
Immune
n=27,085
Cancer
n=27,827
UMAP1
UMAP2
PatientIDs
CS-005
CS-006
CS-007
CS-024
CS-008
CS-009
CS-010
CS-011
CS-11OM
CS-11OV
CS-012
CS-013
CS-014
CS-017
CS-019
D
Clone1
Clone2
Diploid
Immune
CS-024
CS-024
E1
EMT
UMAP1
UMAP2
Clone1 Clone2
latent_time
E1
EMT
S100P
SNAI1
Pseudotime

**Figure S2:**

(**A**) Heatmaps showing RNA-derived copy number variation (rCNV) profiles identify major aneuploid and diploid cell populations. Two distinct clonal populations (Clone 1 and Clone 2) are present within the aneuploid compartment. Heatmap values represent log-transformed copy number alteration signals. The top annotation bar indicates human chromosomes arranged in sequential order.

(**B**) Heatmap displaying the expression of known classical marker genes across CopyKAT-classified cell populations, including predicted cancer cells, non-cancer cells (stromal and immune), and undefined cells.

(**C**) UMAP illustrating cellular heterogeneity across major compartments, including cancer, stromal, and immune populations. Cells are colored by patient of origin, and density contours highlight regions of cell enrichment.

(**D**) UMAP projections of malignant cells from CS-024 with RNA velocity-derived trajectory dynamics overlaid onto inferred clonal identities. Violin plots show latent time distributions across annotated cell states, indicating progression along the inferred trajectory. Line plots depict gene-specific splicing dynamics across pseudotime, with dotted lines for unspliced and solid lines for spliced transcripts; gray shading represents standard deviation.

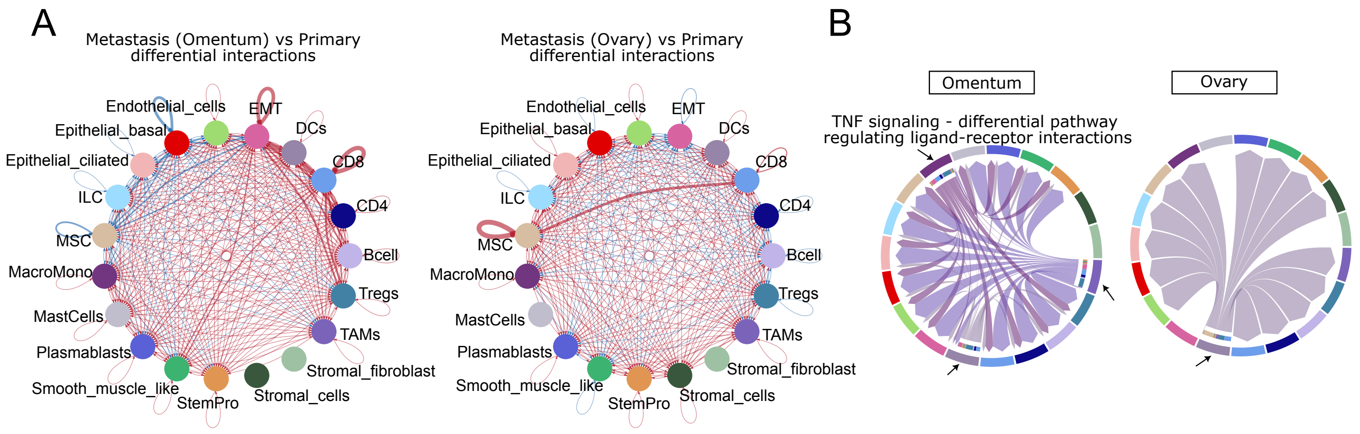
A
Metastasis (Omentum) vs Primary
differential interactions
Metastasis (Ovary) vs Primary
differential interactions
Endothelial_cells
EMT
DCs
Epithelial_basal
Epithelial_ciliated
CD8
ILC
CD4
MSC
Bcell
MacroMono
Tregs
MastCells
TAMs
Plasmablasts
Stromal_fibroblast
Smooth_muscle_like
StemPro
Stromal_cells
B
Omentum
Ovary
TNF signaling - differential pathway
regulating ligand-receptor interactions

**Figure S3:**

(**A**) Interaction diagram illustrating differential cell-cell communication strengths between metastases (omentum and ovary) and the matched primary tumor from patient CS-011, highlighting metastasis-associated remodeling of the tumor microenvironment.

(**B**) Circos plot illustrating the top differentially enriched TNF-α signaling (empirical P-value < 0.05) interactions in CS metastases (omentum and ovary) compared to the matched primary tumor from patient CS-011. The diagram displays both incoming and outgoing signaling contributions for each cell type, with arrows highlighting the most strongly interacting populations, underscoring metastasis-specific remodeling of TNF-α-mediated communication networks. The interaction probability is estimated by analyzing interaction strength of pathway-specific ligand-receptor pairs between cell types.

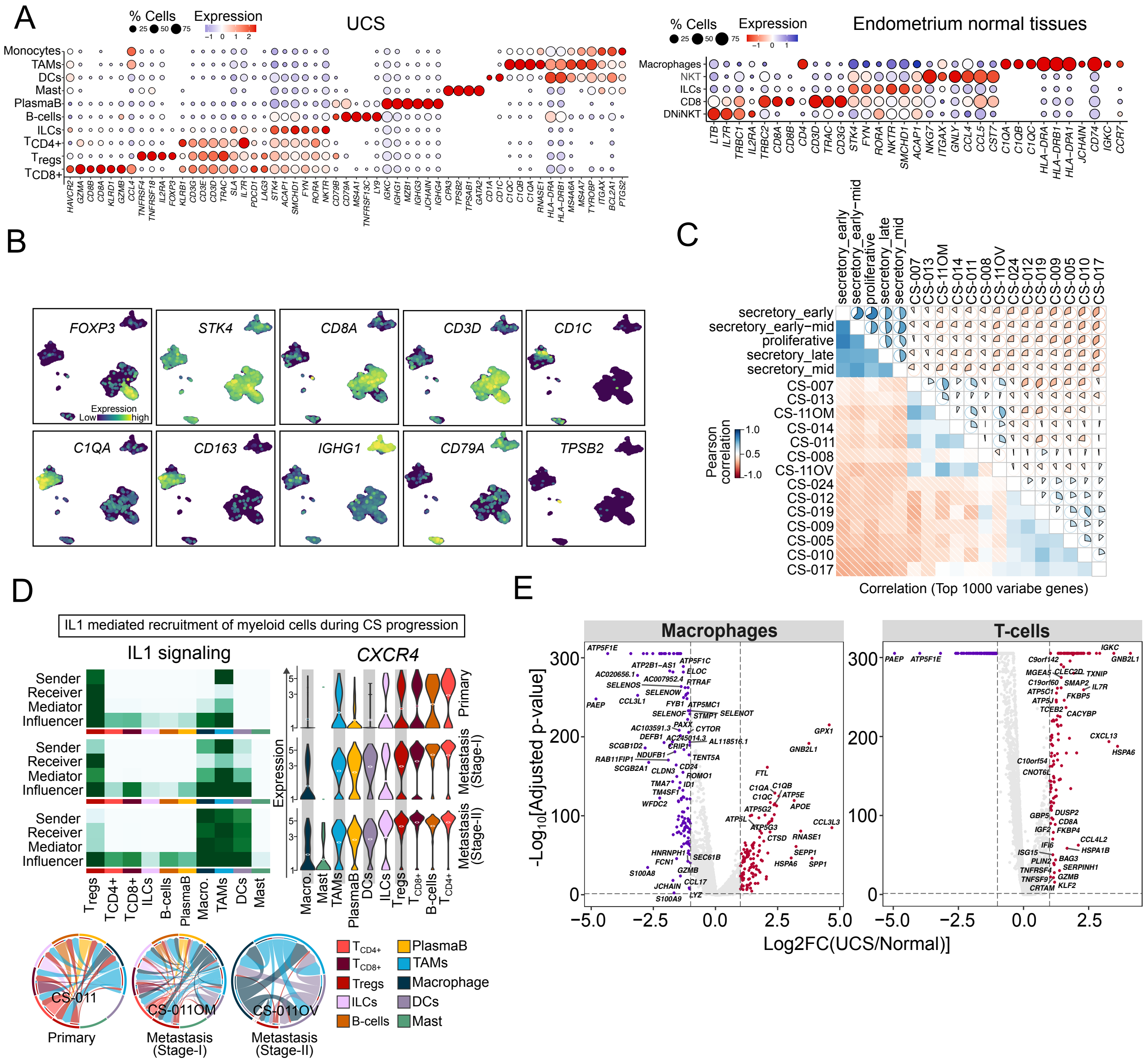
A
% Cells
Expression
UCS
Monocytes
TAMs
DCs
Mast
PlasmaB
B-cells
ILCs
TCD4+
Tregs
TCD8+
HAVCR2 GZMA CD8B CD8A KLRD1 GZMB CCL4 TNFRSF4 TNFRSF18 IL2RA FOXP3 KLRB1 CD3G CD3E CD3D TRAC SLA IL7R PDCD1 LAG3 STK4 ACAP1 SMCHD1 FYN RORA NKTR CD79B CD79A MS4A1 TNFRSF13C LY9 IGKC IGHG1 MZB1 IGHG3 JCHAIN IGHG4 CPA3 TPSB2 TPSAB1 GATA2 CD1A CD1C C1QC C1QB C1QA RNASE1 HLA-DRA HLA-DRB1 MS4A6A MS4A7 TYROBP ITGAX BCL2A1 PTGS2
Endometrium normal tissues
Macrophages
NKT
ILCs
CD8
DNiNKT
LTB IL7R TRBC1 IL2RA TRBC2 CD8A CD8B CD4 CD3D TRAC CD3G STK4 FYN RORA NKTR SMCHD1 ACAP1 NKG7 ITGAX GNLY CCL4 CCL5 CST7 C1QA C1QB C1QC HLA-DRA HLA-DRB1 HLA-DPA1 JCHAIN CD74 IGKC CCR7
B
FOXP3
STK4
CD8A
CD3D
CD1C
C1QA
CD163
IGHG1
CD79A
TPSB2
Expression
Low
high
C
secretory_early
secretory_early-mid
proliferative
secretory_late
secretory_mid
CS-007
CS-013
CS-11OM
CS-014
CS-011
CS-008
CS-11OV
CS-024
CS-012
CS-019
CS-009
CS-005
CS-010
CS-017
Pearson correlation
1.0
0.5
-1.0
Correlation (Top 1000 variabe genes)
D
IL1 mediated recruitment of myeloid cells during CS progression
IL1 signaling
Sender
Receiver
Mediator
Influencer
Tregs TCD4+ TCD8+ ILCs B-cells PlasmaB Macro. TAMs DCs Mast
CXCR4
Expression
Primary
Metastasis (Stage-I)
Metastasis (Stage-II)
Macro. Mast TAMs PlasmaB DCs ILCs Tregs TCD8+ B-cells TCD4+
CS-011
Primary
CS-011OM
Metastasis (Stage-I)
CS-011OV
Metastasis (Stage-II)
TCD4+
TCD8+
Tregs
ILCs
B-cells
PlasmaB
TAMs
Macrophage
DCs
Mast
E
Macrophages
T-cells
-Log10[Adjusted p-value]
Log2FC(UCS/Normal)]
ATP5F1E
PAEP
GPX1
GNB2L1
FTL
C1QA
C1QB
C1QC
APOE
CCL3L3
RNASE1
SEPP1
SPP1
HSPA6
IGKC
CXCL13
IL7R
GZMB
KLF2
CCL4L2
HSPA1B

**Figure S4:**

(**A**) Dot plots displaying cell type–defining marker genes within the immune compartment of UCS tumors (left) and normal endometrium (right). Dot size represents the proportion of cells expressing each gene, while color intensity reflects expression level (red: high; blue: low; white: undetected), enabling comparison of immune cell composition and transcriptional states across conditions. Gene markers are predicted by Seurat with log2fold change ±1 and adjusted p-value < 0.05 determined by Wilcoxon Rank Sum test.

(**B**) UMAP depicting the expression of marker genes across annotated cell types. Light green indicates high expression, whereas dark green reflects low expression, highlighting the spatial distribution of lineage-defining transcriptional programs.

(**C**) Correlation heatmap illustrating transcriptional similarity among individual subjects, including UCS patients and normal endometrium donors, based on the top 1,000 variable genes. Normal donors are stratified by menstrual cycle phase. Blue indicates positive correlation, while red denotes negative correlation, highlighting inter-subject relationships and cycle-associated expression patterns.

(**D**) Heatmap depicting immune cell contributions to IL1 signaling in CS-011 primary and metastatic tumors, with darker green indicating higher activity and lighter green lower activity. Violin plots show *CXCR4* expression across cell types, revealing cell-specific differences between primary and metastatic sites. The circos plot illustrates incoming and outgoing signaling interactions among cell populations, with edge thickness proportional to signaling strength, highlighting rewired communication networks in metastasis. The interaction probability is estimated by analyzing interaction strength of pathway-specific ligand-receptor pairs between cell types.

(**E**) Volcano plots showing differentially expressed genes in macrophages (left) and T cells (right) from UCS tumors relative to normal endometrium. Upregulated genes are shown in red and downregulated genes in blue. Dotted vertical lines indicate the log2 fold-change cutoff (±1), and the horizontal line marks the adjusted p-value threshold (<0.05), highlighting statistically significant transcriptional alterations.

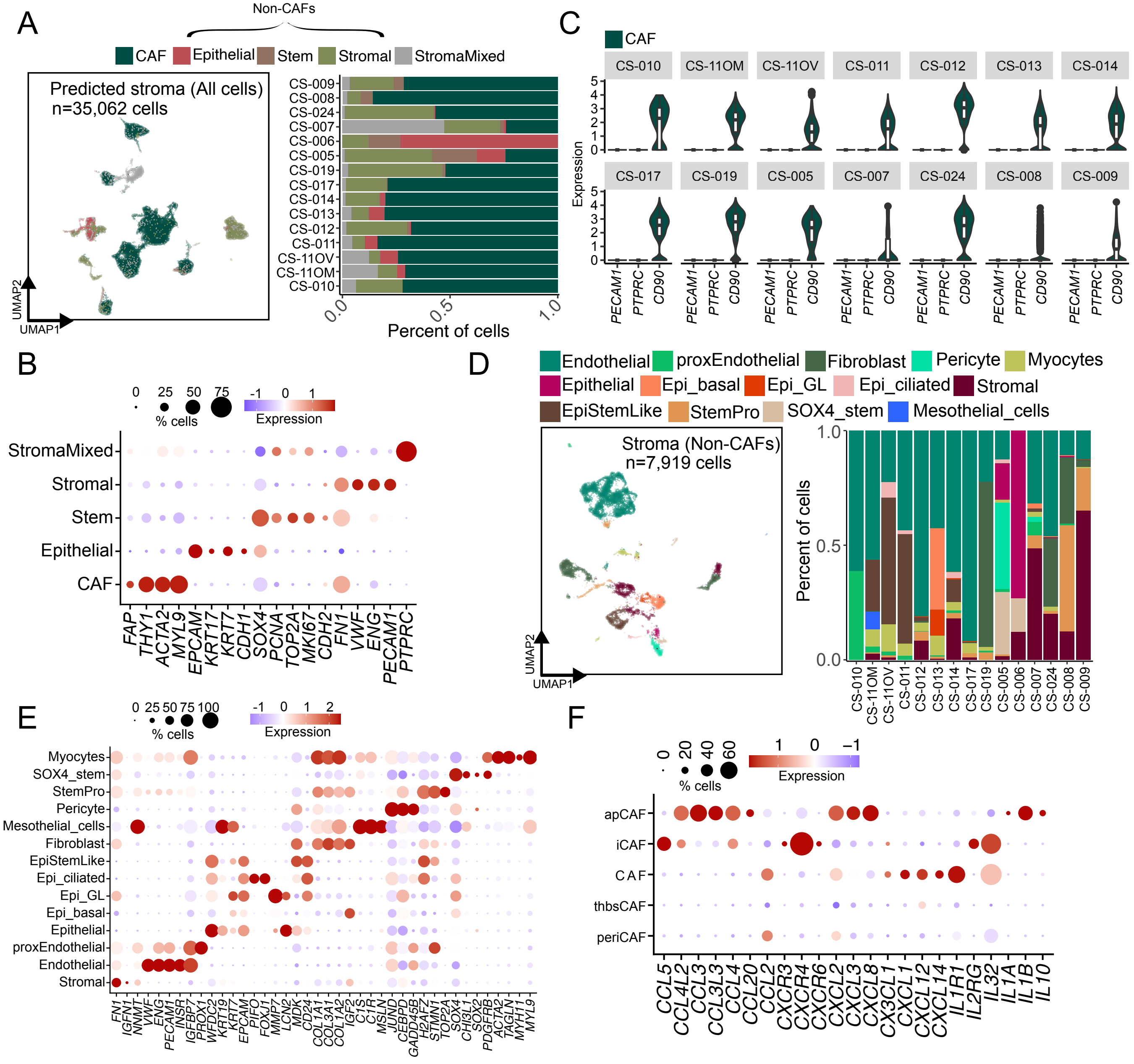
A
Non-CAFs
CAF Epithelial Stem Stromal StromaMixed
Predicted stroma (All cells)
n=35,062 cells
UMAP1
UMAP2
CS-009 CS-008 CS-024 CS-007 CS-006 CS-005 CS-019 CS-017 CS-014 CS-013 CS-012 CS-011 CS-11OV CS-11OM CS-010
Percent of cells
C
CAF
CS-010 CS-11OM CS-11OV CS-011 CS-012 CS-013 CS-014
CS-017 CS-019 CS-005 CS-007 CS-024 CS-008 CS-009
Expression
PECAM1 PTPRC CD90
B
% cells
Expression
StromaMixed Stromal Stem Epithelial CAF
FAP THY1 ACTA2 MYL9 EPCAM KRT17 KRT7 CDH1 SOX4 PCNA TOP2A MKI67 CDH2 FN1 VWF ENG PECAM1 PTPRC
D
Endothelial proxEndothelial Fibroblast Pericyte Myocytes
Epithelial Epi_basal Epi_GL Epi_ciliated Stromal
EpiStemLike StemPro SOX4_stem Mesothelial_cells
Stroma (Non-CAFs)
n=7,919 cells
Percent of cells
E
% cells
Expression
Myocytes SOX4_stem StemPro Pericyte Mesothelial_cells Fibroblast EpiStemLike Epi_ciliated Epi_GL Epi_basal Epithelial proxEndothelial Endothelial Stromal
FN1 IGFN1 NNMT VWF ENG PECAM1 INSR IGFBP7 PROX1 WFDC2 KRT19 KRT7 EPCAM PIFO FOXJ1 MMP7 LCN2 MDK CD24 COL1A1 COL3A1 COL1A2 IGF2 C1S C1R MSLN JUND CEBPD GADD45B H2AFZ STMN1 TOP2A SOX4 CHI3L1 SOX2 PDGFRB ACTA2 TAGLN MYH11 MYL9
F
% cells
Expression
apCAF iCAF CAF thbsCAF periCAF
CCL5 CCL4L2 CCL3 CCL3L3 CCL4 CCL20 CCL2 CXCR3 CXCR4 CXCR6 CXCL2 CXCL3 CXCL8 CX3CL1 CXCL1 CXCL12 CXCL14 IL1R1 IL2RG IL32 IL1A IL1B IL10

**Figure S5:**

(**A**) UMAP and accompanying bar graph illustrating the major stromal cell populations, including CAFs and non-CAFs (epithelial, stromal, and stem cells), and their distribution across patients, highlighting interpatient heterogeneity in stromal composition.

(**B**) Dot plots displaying cell type-defining marker genes within the stromal compartment of CS tumors. Dot size represents the proportion of cells expressing each gene, while color intensity reflects expression level (red: high; blue: low; white: undetected), supporting annotation of stromal subpopulations. Gene markers are predicted by Seurat with log2fold change ±1 and adjusted p-value < 0.05 determined by Wilcoxon Rank Sum test.

(**C**) Violin plots showing the expression of *PTPRC*, *PECAM1*, and *CD90* across CAF populations from individual UCS patients, highlighting patient-specific variability and confirming lineage-associated marker profiles.

(**D**) UMAP and accompanying bar graph illustrating the identified non-CAF subtypes (epithelial, stromal, and stem cells) and their distribution across patients, revealing interpatient variability in non-CAF composition.

(**E**) Dot plots displaying cell type-defining marker genes across non-CAF populations in UCS tumors. Dot size represents the proportion of expressing cells, and color intensity reflects expression level (red: high; blue: low; white: undetected), supporting precise annotation of non-CAF subtypes. Gene markers are predicted by Seurat with log2fold change ±1 and adjusted p-value < 0.05 determined by Wilcoxon Rank Sum test.

(**F**) Dot plot illustrating the expression of chemokines and their receptors across CAF subtypes, with dot size indicating the percentage of expressing cells and color intensity denoting expression level, highlighting subtype-specific signaling potential.
